# Controlling for temporal discounting shifts rats from geometric to human-like arithmetic bisection

**DOI:** 10.1101/245092

**Authors:** Charles D Kopec, Carlos D Brody

## Abstract

How our brains measure the passage of time is still largely open for debate. One behavioral task commonly used to study how durations are perceived is the Temporal Bisection Task, in which subjects categorize time durations as either “short” or “long.” The duration equally likely to be categorized as short or long is known as the bisection point. It has been consistently demonstrated that for humans, the bisection point is near the arithmetic mean of the longest and shortest durations the subject was trained on. In contrast, for non-human subjects it has been consistently found near the geometric mean. This difference implies that humans may process or represent temporal durations differently than other species. Here we present a behavioral model that reconciles the differences by demonstrating that rats’ performance on this task is driven not only by their noisy estimates of duration, but also by the temporally-discounted value of future rewards. The model correctly predicts shifts in the bisection point induced by unequal rewards and explains otherwise-paradoxical psychometric reversals documented three decades ago. Furthermore, as predicted by the model, we found that modifying the Temporal Bisection Task to eliminate the temporally-discounted reward component shifted the rats’ bisection point from the geometric mean to the arithmetic mean, thus bringing the rat results into line with the human results. We therefore propose that humans and rats (and perhaps other non-human subjects as well) process temporal information similarly, and that the difference between them in the Temporal Bisection Task may be simply due to rats weighing temporal discounting of future rewards more strongly than humans.

## Introduction

The neural mechanisms by which we perceive and measure the passage of time largely remain a mystery [1–7]. For the past century researchers have developed behavioral paradigms to address questions such as: Is time in our brain represented linearly like distance, logarithmically like pitch, or qualitatively like color? [8–10]; How does our emotional state affect the perceived passage of time? [11–13]; How do we compare durations? [1,14–16]; and, Can memories of durations be influenced? [17–20]; just to name a few.

The Temporal Bisection Task is designed to address how one innately categorizes durations [14,21]. A subject is first trained to discriminate two durations, called the long and short reference durations, after which a series of intermediate probe durations are introduced. The subject is then instructed, for each probe duration, to indicate which reference duration they estimate it is most similar to. Here feedback is only given for the reference durations so as to not influence the subject’s categorization of the intermediate probe durations. The probe duration they report with equal probability as being most similar to “long” or “short” is referred to as the point of subjective equality, or bisection point. The position of this categorization boundary offers clues as to how time is represented and how durations are compared mentally [14,16,22,23].

Since its first application to humans almost two and a half decades ago [14,24], numerous groups have found that we tend to bisect near the arithmetic mean of the reference durations ([14,25–28], see [16] for review and explanation for why human subjects occasionally appear to bisect near the geometric mean). However, for almost four decades a range of non-human species have been trained on this task and consistently been found to bisect near the geometric mean of the reference durations [21,29–35]. While this difference may seem trivial, it potentially means that non-human species may perceive or process temporal information differently from us. For example, humans may perceive time linearly, while animals may perceive time logarithmically [10] (although see [8,9] for evidence arguing against the logarithmic hypothesis). Or animals may compare durations using ratios [22,36,37] while humans use differences [14,16]. Such fundamental differences would call into question what we can learn about human time perception from studying non-human subjects.

Here we present a behavioral model capable of reconciling rat and human behavior on the Temporal Bisection Task. The model brings together two well-known temporal properties that have nevertheless usually been treated separately. First, noisy *scalar timing* in the estimate of when the reward will become available, which refers to the widespread observation that variability in the estimate of a temporal duration scales proportionately with the duration [1,8,14,22,36,38–43]. Second, *temporal discounting* of the value of a reward, which refers to the separate but also widespread observation that rewards that are distal in time are valued less than rewards that are proximal in time [44–54]. The model uses these two aspects as drivers of response behavior. The bisection point is then reinterpreted not as the point where the duration *percept* is equally similar to the two references, but instead as the point when the two motor response options, “short” and “long”, have equal average *value* to the subject.

With this model we are able to demonstrate that behavior on a single duration estimation task, the Peak Interval Task, which does not involve classification of different durations, is nevertheless able to predict geometric bisection on the Temporal Bisection Task. The prediction suggests that geometric bisection may be strongly dependent on differential temporal discounting of reward for the two references. Consistent with the idea that reward value is an important force driving response behavior in timing tasks, the model makes successful quantitative predictions as to how the bisection point should shift if the rewards are unbalanced between the two response options. The model also offers a novel explanation for the decades-old and otherwise puzzling observation in non-human subjects, that as probe durations increase beyond the reference “long” duration, the fraction of “long” responses begins to decrease rather than continuing to increase [32,33,35].

Our model suggests that in order to assess a rat’s true *perceptual* bisection point, defined as the duration they perceive as equidistant between the two trained reference durations, we should modify the Temporal Bisection Task to eliminate temporally-discounted reward value as a driver of response behavior. We found that this modification caused rats to shift their bisection point from the geometric to the arithmetic mean, in line with human performance. This result, eliminating the difference in bisection points between human and non-human subjects, suggests that human and non-human subjects may process temporal information similarly, which in turn justifies investigation of the neural basis of time perception in non-human subjects as a means towards elucidation of neural mechanisms of human time perception. We suggest that the difference between humans and non-humans in bisection point in the traditional Temporal Bisection Task may simply be a matter of different strategies in their performance of the task, not a fundamental difference in their time perception mechanisms.

## Results

In the Temporal Bisection Task, subjects are first trained to discriminate two reference durations: “short” and “long”. Upon achieving discrimination at some high degree of accuracy, intermediate length probe durations are introduced. Human subjects are instructed to report which reference duration they estimate the current stimulus is most similar to. The bisection point is computed as the duration the subject is equally likely to classify as “long” or “short”, and has traditionally been interpreted as the duration that the subject perceives as equidistant between the two reference durations [14,21].

Here we trained 10 rats on a modified version of the Temporal Bisection Task. Trials started by turning on an LED in the center port, in response to which trained rats poked into the center port, marking time t=0 (Figure 1A). This poke also initiated an auditory tone that terminated after either a short duration T_short_ or after a long duration T_long_. If the tone was of duration T_short_, a poke after T_short_ into the left port (which we called the “short” port) elicited a water drop reward and terminated the trial. In contrast, a poke into the right port (which we called the “long” port) elicited 8 seconds of white noise in place of the reward. If the tone was of duration T_long_, the contingencies were reversed. Any pokes before the end of the stimulus, on both “short” and “long” trials, had no effect but were monitored and are used in our analysis and model. Typical behavior on “long” trials was for rats to initially poke repeatedly into the “short” port, until time T_short_ has elapsed; some time after T_short_ they moved to the “long” port and poked there until T_long_ (Figure 1B,E). We trained rats on four different (T_short_, T_long_) pairs, in each pair keeping T_long_ at ten times T_short_ (i.e. *T_long_ =* 10*T_short;_* pair 10.39s, 3.9s; Pair 2 0.78s, 7.8s; Pair 3 1.57s, 15.7s; and Pair 4 3.14s, 31.4s see Methods).

**Figure 1.**
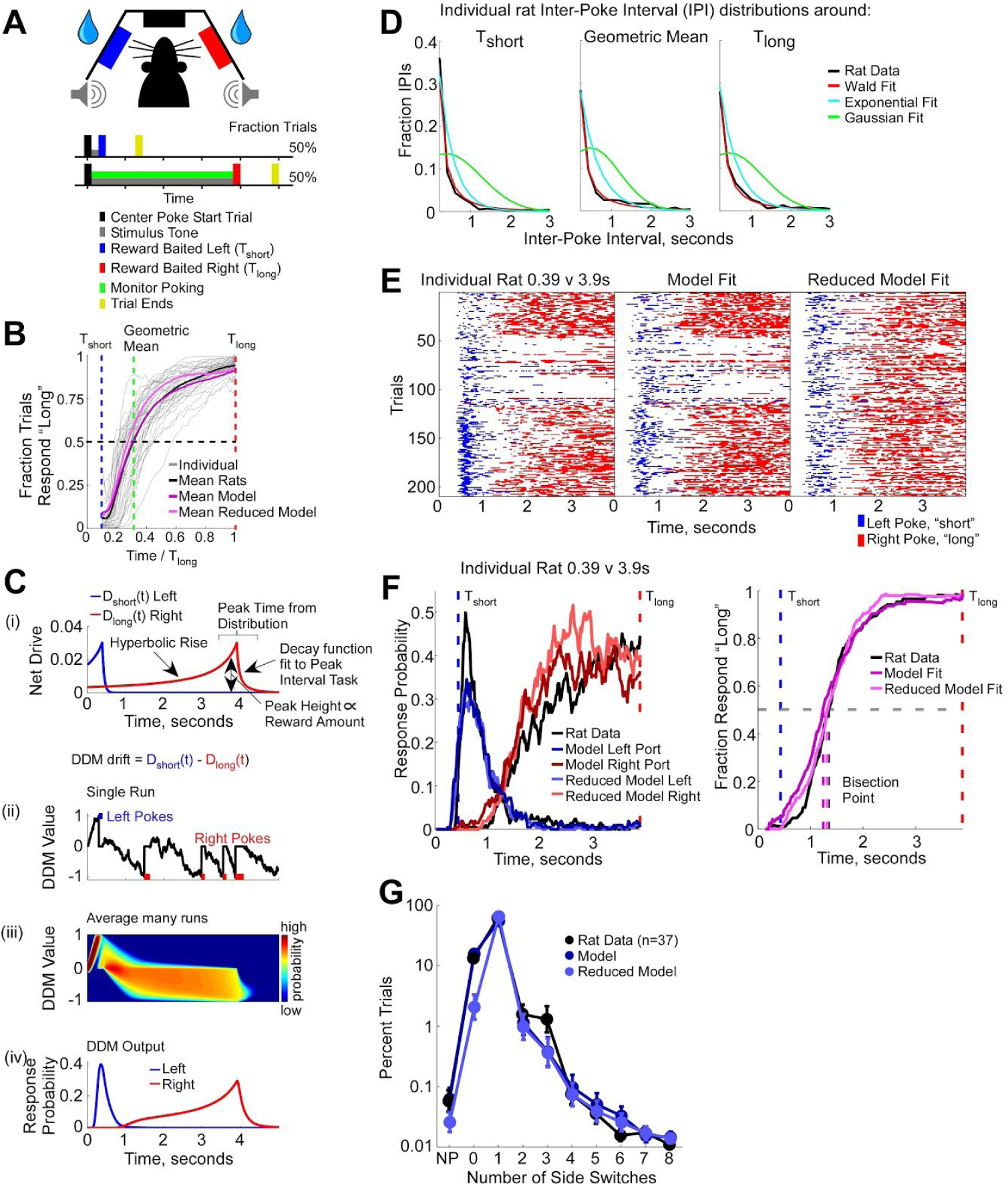
Performance and modeling of the Temporal Bisection Task. **A)** Upper: diagram of the behavioral chamber containing 3 nose ports, stereo speakers, and drink tubes in the left and right ports for reward delivery. Lower: Temporal Bisection Task structure. On short tone trials, rats are rewarded for poking left; on long tone trials, they are rewarded for poking right. Nose pokes during the long tone are monitored and analyzed. **B)** Analysis of “long” trials. Each line represents the fraction of trials in which a rat was at the “long” (right) nose port, as a function of time between T_short_ and T_long_. The bisection point is defined here as the time when the subject is equally likely to be at the “short” or “long” port. C) Behavior model: The difference between a “net drive” function for each of the two response ports is used as the drift term in a drift-diffusion model (DDM). In this DDM, reaching the upper (or lower) boundary corresponds to a initiating a left (or right) poke. (i) Example net drive functions D_short_(t) and D_long_ (t) associated with generating “short” and “long” responses in the left and right nose ports respectively. (ii) Single run of the DDM using the net drive functions in (i) with the left and right nose pokes produced. (iii) Fokker-Planck solution for the DDM showing how the full probability distribution over DDM values evolves over time. This is equivalent to an infinite number of runs of the DDM using the net drive functions in (i). (iv) Probability of producing a “short” (left) or “long” (right) response taken from DDM threshold crossing in (iii). **D)** In agreement with predictions of a DDM process, inter-poke intervals are well fit by a Wald distribution. The panels show histograms of the inter-poke intervals (IPIs) taken from three example individual experiments: rat K006 performing 1.57 v 15.7s, IPIs from a 2s window around T_short_; rat K014 performing 0.78 v 7.8s, IPIs from a 2s window around the geometric mean; rat K008 performing 0.39 v 3.9s, IPIs from a 2s window around T_long_. Each panel shows the best fit Wald, exponential, and Gaussian functions. **E)** Left: Poke rasters across all trials from an individual rat performing the Temporal Bisection Task with reference durations T_short_ = 0.39s and T_long_ = 3.9s. Center: Full model produced poke rasters. Right: Reduced model produced poke rasters. **F)** Left: Comparison of rat (black) and model fit (color) average “short” (left) and “long” (right) response probability. Right: Comparison of rat and model fit psychometric curves, computed by interpolating the poke rasters in **E**, see methods. G) Percent of long trials on which various numbers of side switches are produced. NP: no pokes produced before T_long_. Note the logarithmic scale on the vertical axes. Error bars are S.E.M.

Instead of using a particular set of probe duration stimuli, as is classically done in this task [21], we used an approach previously taken with pigeons [21,29,30] to measure the subject’s response to intermediate duration stimuli. We simply monitored poking during the long stimulus trials. We assume that if a stimulus were to terminate at an intermediate duration, the rat’s report for that stimulus would correspond to whichever port the rat was poking into at that time. Thus, at each point in the time interval T_short_<t<T_long_, we assess the fraction of trials in which the rat was in the “short” port and the fraction of trials the rat was in the “long” port, and take that as the fraction of “short” and “long” reports that the rat would have made for a stimulus of duration t (see Methods). As previously found with pigeons [21,29,30,32], this modified version of the Temporal Bisection Task replicated results from the standard version of the task, including a sigmoid response as a function of t (Figure 1B) and a bisection point at the geometric mean of T_short_ and T_long_ [21,22,29,30,32]. In addition, this approach offers us, on each trial, a continuous moment-by-moment response function throughout the interval T_short_<t<T_long_, rather than just a single response at a single duration.

We first sought to develop a unified modeling framework for describing real time moment-by-moment response behavior across a range of temporal processing tasks. Our modeling framework relies on three concepts. First, at any point in time, and for each relevant response, there is a number that represents the net subjective value, or incentive for the subject, to elicit that response. We refer to this function as the “net drive” (Figure 1C i). Second, the net drive for a response acts as the input to a noisy decision process, instantiated as a drift diffusion model (DDM, [55]). The DDM integrates the net drive function(s) D(t) and actual responses are initiated when the DDM exceeds a threshold (Figure 1C ii, see [56,57] for examples of using DDMs in value-based decision-making). The greater the net drive, the more likely and the more frequent the response; but the decision process itself is stochastic. Third, when the subject has a choice of two responses, but can only elicit one at a time, the drives for the two responses compete by moving the DDM towards opposite bounds (Figure 1C). Reaching a bound corresponds to initiating a response poke.

The DDM value V(t) evolves according to the following equation

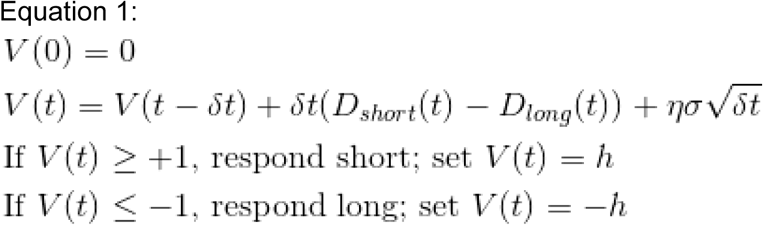

where D_short_(t) represents the net drive for producing a “short” response, D_long_(t) is the net drive for producing a “long” response, δt is the time step size, σ is the diffusion noise constant, η is drawn from a normal distribution with zero mean and unit standard deviation, and h is a hysteresis parameter controlling the value V(t) is set to following a response.

One prediction of a DDM, given constant drift and Gaussian diffusive noise, is that the time to bound crossing is described by a Wald distribution [58–60]. In our data this is analogous to the inter-response interval (the time between nose pokes, IPI, inter-poke interval). Supporting our use of a DDM, histograms of the inter-poke intervals taken at fixed times (around T_short_, T_long_, or the geometric mean of T_short_ and T_long_) during repeated identical trials are better fit by a Wald distribution than either Gaussian or Exponential distributions (see Figure 1D for example IPI histograms taken from 3 different rats on the Temporal Bisection Task, overall across all rats and time periods tested within the trials a Wald distribution fits the data better than a Gaussian or an exponential, p<<0.01).

The form of the “net drive” function, D(t) in equation 1, itself directly influences the profile of response behavior. We began from the concept of temporally-discounted reward, which states that the value of a reward is inversely proportional to the delay to that reward [44,45], implying that future rewards are valued less than immediate rewards. Across a range of species, it has been demonstrated that future rewards are discounted following a hyperbolic function [44,45,47,53,54,61] (Supp Figure 2C, see methods). Therefore, as a reward approaches in time within a trial, its value should rise hyperbolically. The height to which the net drive function rises is proportional to the absolute magnitude of the reward, and the time at which the function peaks is given as the subject’s noisy estimate of the reward time on that trial (Figure 1C i). Following the estimated time of a reward, response rates decrease [62], therefore the net drive function should similarly decay. The form of the decay function used here was fit from data on the Peak Interval Task (Supp Figure 4, see methods) where subjects only have to estimate a single duration. The net drive function we used was therefore given by the following.

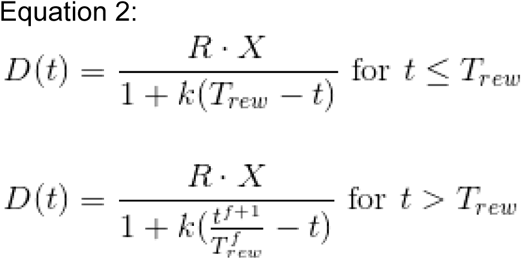

The model parameters are as follows: R is a scaling constant, X is the number of drops of reward offered if this is the correct response, k is the hyperbolic discounting constant, T_rew_ is the subject’s noisy estimate of the reward time (T_short_ or T_long_), t is the actual time within a trial, and f is a constant that controls the steepness of the decay following an omitted reward.

The modeling framework as described in Equations 1 and 2 is not specifically designed to solve any particular temporal processing task. It merely describes how the incentive value of producing a task-relevant response varies as a function of time, and produces those responses in a stochastic manner. The model parameters are fit to data by maximizing the likelihood of the model producing the experimentally-observed moment-by-moment response behavior. Below we will demonstrate how this model can not only fit response behavior on the Temporal Bisection Task, but also provide a novel insight as to the strategy non-human subjects may be using to solve the task. For a complete description of the model and examples of how it can fit moment-by-moment response behavior on two additional temporal processing tasks, the Delay Discounting Task and the Peak Interval Task, please see the supplemental text and Supplemental Figures 1-5.

To apply the modeling framework to the Temporal Bisection Task we defined two net drive functions (Figure 1C i). D_short_(t) rises hyperbolically to the subject’s noisy estimate of T_short_, decaying afterwards, and pulls the DDM towards the bound associated with making responses in the “short” nose port. D_long_ (t) rises following a hyperbolic function with the same curvature as D_short_(t) but now peaking at the subject’s noisy estimate of T_long_, decaying afterwards, and pulls the DDM in the opposite direction towards the bound associated with making responses in the “long” nose port (Figure 1C ii).

Given a set of model parameters, we evolve the full probability distribution of the DDM (Figure 1C iii) and compute the probability of the model producing a “short” or “long” response at each moment in time (Figure 1C iv). We fit the model parameters to data from each experiment independently, maximizing the likelihood of the model producing the moment-by-moment response behavior on “long” trials for each rat on each T_short_ T_long_ duration pair (see methods).

Based on the time estimation literature [1,8,14,22,36,38–43] we know the subject’s estimate of T_short_ and T_long_ will be subject to noise. Behavioral models typically account for this noise by describing the estimate of a duration as a distribution from which a unique value is pulled on each trial [14,39]. Translating this back to the subject, the rat should therefore have a discrete and unique value for T_short_ and T_long_ on each trial and behave accordingly. For example, on trial 10 the subject’s estimate of T_short_ may be 2.5s and for T_long_ may be 6s while on trial 11 they may be 1.7s and 5.1s respectively. Here we are fitting our model to the moment-by-moment trial-by-trial poking behavior (Figure 1E) rather than the summary choice statistics (did he earn the small or large reward) or the average response profile (Figure 1F). In order to determine the model parameters that maximize the likelihood of replicating the exact nose-poking pattern produced by the rats for each trial, we must know the value that was pulled for T_short_ and T_long_ on each trial. It is not enough to know that the estimate of T_short_ follows a certain distribution with a specific mean and standard deviation, since the trials the model performs would not correspond to the trials the subject performs. Following the example above, the model by chance may pull a value of 4s for T_short_ on trial 10 causing it to produce a different response pattern from the subject, leading to a poor fit, but not because the model is necessarily bad but because the stocasticity of the model and the rat are not in sync. Therefore, since the subject likely draws a discrete value from their noisy estimate for T_short_ and T_long_ on each trial, we found those values that maximize the log-likelihood of the model replicating the subject’s response behavior (see Methods). In doing so we not only avoid making an assumption as to what type of distribution should describe the subject’s estimate (Gaussian, log-normal, gamma…) but also can directly assess that distribution by simply plotting the histogram of the best fit values for T_short_ and T_long_ across trials. We found that the model estimates for T_short_ and T_long_ on each trial form distributions that, when scaled by the actual reward time, superimpose well on each other (Supp Figure 7) consistent with scalar timing [36,41].

By maximizing the likelihood of the model replicating the precise poking behavior produced by the subject in a moment-by-moment and trial-by-trial manner (at 20ms resolution for both the left and right nose ports, see Methods), rather than simply their final “choice”, we are attempting to fit 100 data points per second per trial that the subject performs. Finding the best fit values for T_short_ and T_long_ on each trial therefore does not lead to over-fitting as the number of fit parameters per experiment is always orders of magnitude below the number of data points we are attempting to fit.

However, once we find the best-fit parameters for the model including the values for T_short_ and T_long_ on each trial (which we refer to as the “full” model) we then attempted to reduce the model parameters by describing the distribution of T_short_ and T_long_ as a 2-parameter log-Normal function:

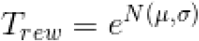

where N is a normal distribution with µ mean and σ standard deviation.

While this “reduced” model is no longer in sync with the rat’s performance on a trial-by-trial basis (Figure 1E) it is still able to capture the overall response behavior well (Figure 1B,F,G) with a substantially reduced set of free parameters. While the reduced model’s psychometric curve (Figure 1B) is shifted slightly to the left (earlier in time) it on average produces a bisection point normalized by T_long_ at 0.298 +/- 0.022 (mean +/- standard error) which is not statistically different from the full model’s normalized bisection point (0.301 +/- 0.015, p=0.23), the rat’s normalized bisection point (0.304 +/- 0.014, p=0.36) or the normalized geometric mean of T_short_ and T_long_ 0.316 (p=0.40).

One example of a rat’s response behavior, average response probability, and psychometric curve alongside the model fit is shown in Figure 1E,F (see Supp Figure 6 for all individual rat data and model fits). The model not only does a good job of fitting the overall response behavior, psychometric curve, and more specifically bisection at the geometric mean across all rats (Figure 1B), but is also able to account for atypical response behavior that occurs on a minority of trials. While it is typical for a rat to display one “change of mind” on a “long” trial, switching from the “short” port to the “long” port on average around the geometric mean, occasionally rats produce many more side switches (3.0% of trials) or no side switches (13.3%), occasionally making no responses at all (0.06%, Figure 1G, see [30] for similar behavioral finding). Such behavior has previously been interpreted as a subject’s inattention to the task at hand [63], and occasionally excluded from analysis [64]. Wth this modeling framework, the stochastic component of the decision process allows such trials to exist and the model overall is able to fit the frequency of such atypical trials well (Figure 1G).

Our framework makes several predictions, each of which we discuss and confirm below. First, we reasoned that since geometric bisection in the Temporal Bisection task was successfully modeled as based on two competing but otherwise independent single-duration drives (Figure 1), it follows that fitting the model to a single-duration estimation task would produce parameters that would themselves predict geometric bisection. Second, reward is an integral force driving behavior in the model; the model should therefore be able to quantitatively predict the behavioral effects of varying reward magnitudes. Third, the net drive in the model decays after the expected reward baiting time. This should allow the model to account for psychometric “reversals” at long probe times, i.e., the observation in the Temporal Bisection Task that as probe durations increase beyond the “long” reference duration, the fraction of “long” responses begins to decrease rather than continuing to increase. We now describe these in turn.

We first asked whether parameters obtained from behavior on a single-duration estimation task could predict geometric bisection on a duration classification task. Others have also suggested that performance on the bisection task may be seen as a competition between two fixed interval schedules [65]. Here we sought to empirically test this prediction. We began by training rats on the Peak Interval Task [62]. a classic single-duration estimation task. In this task, rats start a trial by nose poking into the center port; this event marks time t=0. On the majority of trials (randomly chosen 85% of trials), a reward will be baited in the left nose port after a fixed duration. On the remaining 15% of trials no reward is baited and response behavior is monitored (Supp Figure 4). In these 15% of trials, rats typically nose poke in the reward port with a frequency that increases before the baiting duration, and decreases thereafter. The time of peak nose poking frequency is traditionally taken to indicate the rats’ estimate of the baiting duration. We analysed these reward-omitted trials by fitting our modeling framework to the rats’ response behavior, using a single net drive function of the same form as described above. We found that the model can provide a good fit to the rats’ moment-by-moment response behavior in this task (Supp Figures 4,5). We then took the best-fitting model parameters values from the rats’ behavior on the Peak Interval Task, and used those parameter values to run the model, now with competing drives so as to model the Temporal Bisection Task. We found that the predicted bisection point was indistinguishable from the geometric mean (see Figure 2A, mean T_bisection_/T_long_ = 0.280 +/- 0.040, p = 0.351 that T_bisection_/T_long_ is significantly different from the geometric mean value 0.316; in our experiments we used T_long_ = 10x T_short_). Thus behavior on the Peak Interval Task, which involves no temporal classification, predicts geometric bisection on a duration classification task, Temporal Bisection.

**Figure 2.**
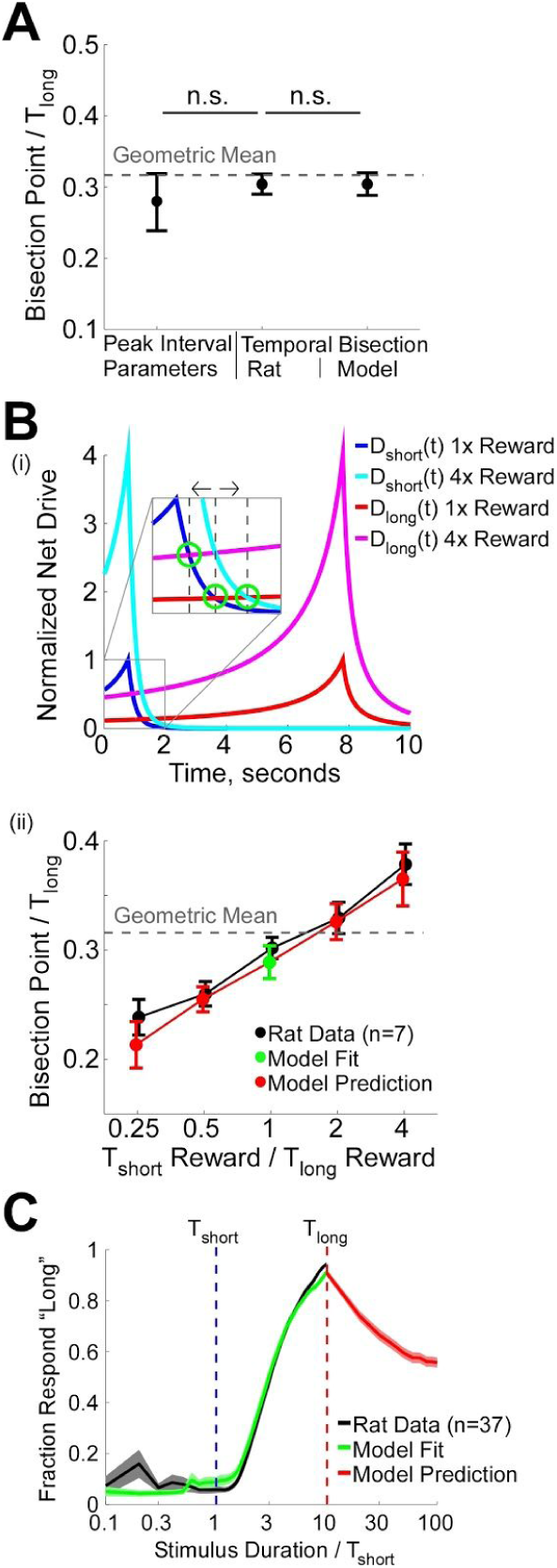
Testing model predictions. **A)** Fitting the model to data from a single-duration estimation task produces parameter values that predict geometric bisection in the Temporal Bisection Task. Left data point: Parameter values fit to rat data on the single-duration Peak Interval Task predict a bisection point indistinguishable from geometric bisection in the Temporal Bisection Task. Middle: Bisection point from rat behavioral data in the Temporal Bisection Task, shown for comparison. Right: Parameters fit to data from the Temporal Bisection Task, also shown for comparison. n.s. not significant (p>0.05). **B)** The modeling framework quantitatively predicts shifts in the bisection point induced by unequal rewards at the “short” and “long” ports. (i) Schematic illustrating how the modeling framework predicts shifts in the bisection point following varying reward magnitude. Net drive functions D_short_(t) and D_long_(t) associated with T_short_ and T_long_ durations respectively are shown for 1x and 4x reward conditions. The inset highlights the shift in the intersection of the two net drive functions. Green circles indicate places around intersections referenced in the text, (ii) Mean bisection point as a function of the ratio of the rewards offered following T_short_ and T_long_. The model was fit to the balanced reward condition only (green center point), and the resulting parameter values when run on the unbalanced reward conditions (red) are in good agreement with the rats’ measured bisection points (black). **C)** The model predicts a non-monotonic psychometric function with a decrease in the fraction of “long” responses following T_long_. Average rat data (black) and model fit (green) shown for the period t<T_long_. Average model prediction (red) generated using the best fit model parameters for each individual experiment from the period t<=T_long_ run on durations T_long_<t<=10xT_long_. Our rats do not show a significant psychometric reversal in the brief period before T_short_, a finding the model can be well fit to. Note the logarithmic scale of the horizontal axis ranging from 1 /10^th^ T_short_ to 10xT_long_. Error bars S.E.M. across individual experiments (each rat performing the task given a pair of reference durations constitutes an experiment).

Next we tested the model’s predictions regarding reward sensitivity. According to our modeling framework, reward is an integral force driving response behavior, and varying the amount of reward offered following either the “short” or “long” reference duration on the Temporal Bisection Task produce quantitative shifts in the bisection point that are precisely predicted by the model. For example, if 4 drops of water were offered following a correct “short” response (X=4 in equation 2), the height of the net drive function associated with producing “short” responses, D_short_(t), would increase by an equivalent factor. This would delay when the net drive for both responses were equivalent, and thereby delay the bisection point (Figure 2B i, intersection of cyan and red lines). Increasing the reward associated with a correct “long” response would have the opposite effect: a greater height of the net drive function associated with producing a “long” response, D_long_ (t), would hasten when the net drive functions for both responses are equivalent (Figure 2B i, intersection of blue and magenta lines), thereby hastening the bisection point. (Note that according to our modeling framework, the bisection point in the Temporal Bisection Task is related to the intersection of the D_short_(t) and D_long_ (t) net drive functions described by equation 2, but this intersection point and the bisection point may not be identical, since bisection is calculated from poke rasters which are produced by passing the net drive functions through the DDM of equation 1.)

To test this prediction we trained rats on the Temporal Bisection Task using a single T_short_ T_long_ pair (0.78 and 7.8s respectively) but varied the reward ratio across blocks of sessions offered between the two response options (“short” reward : “long” reward) across a 16-fold range (1:4, 1:2, 1:1, 2:1, 4:1, see methods). Reward amounts were kept fixed across sessions until all data for that reward pair was collected. Upon transition to a new reward amount performance had to reach 80% correct for an entire session before data collection began. Reward amounts were tested in random order for each rat. We fit the model parameters to performance on trials with the balanced reward condition only (1:1) and then used those parameters to run the model on the unbalanced reward conditions. The model not only accounted for the qualitative shift in the bisection point, delaying when a “short” response was rewarded more and hastening when a “long” response was rewarded more, but predicted the precise magnitude of the shift as well (Figure 2B ii).

Next we turned to an old, and largely unexplained observation [23] of “psychometric reversals” for probe durations longer than the “long” reference [32,33,35]. The data led authors of these studies to propose, as we do, that non-human subjects do not classify durations simply based on the traditionally assumed perceptual similarity strategy. Siegel [32] trained rats on a classic version of the Temporal Bisection Task but tested them with probe durations both within and beyond the range of T_short_<t<T_long_. If rats were classifying the probe durations according to which reference duration they perceived it as most similar to (the Similarity Rule [22]), one would predict a monotonic psychometric function, meaning durations longer than T_long_ would be as much or more likely to be classified as “long” than the T_long_ duration, and durations shorter than T_short_ would be classified as “short” as much or more than the T_short_ duration. However, Siegel found that rats produced a non-monotonic psychometric function [32], reducing the probability of reporting probes t>T_long_ as most similar to T_long_. This finding was later and again recently replicated in pigeons [33,35].

We found that our modeling framework predicts such a psychometric reversal. The difference between the net drive functions D_short_(t) and D_long_(t) is maximal at the subject’s estimate of the reference durations (Figure 1C i). The model is therefore maximally biased to produce “short” (“long”) responses at its estimate of T_short_ (T_long_). After T_long_, the value of D_short_(t) and D_long_(t) rapidly approach one another. Therefore, the bias to produce a “long” response over “short” is reduced, resulting in a reversal for t > T_long_. However, because the hyperbolic rise of the net drive function before a reward is more gradual than the decay afterwards (equation 2), and because one can only probe a short range of durations prior to T_short_, our modeling framework predicts no significant reversal in the period t < T_short_. Findings of reversals prior to T_short_ are inconsistent, but when found (see [32] experiment 2 and [35] type 2 trials) the magnitude of the reversal tends to be small. In the brief period between the start of the trial and T_short_, our rats do not produce a significant reversal in their response preference (Figure 2C, black), a finding that our model can be well fit to (Figure 2C, green). Using the best fit model parameters however, we predict a robust psychometric reversal for durations significantly greater than T_long_, with response preference slowly asymptoting towards chance (Figure 2C, red), consistent with previous experimental findings [32,33,35].

To summarize what has been presented so far, we found that behavior in the Temporal Bisection Task was well described with two competing net drive functions that each had a form derived from considerations that do not involve duration classification: the net drives have a hyperbolic rise to the reward time (as prescribed by temporal reward discounting), and a decay following an omitted reward obtained from fits to a single-duration estimation task (the Peak Interval Task). With these net drives, the framework successfully accounted for geometric bisection and for moment-by-moment response behavior in the Temporal Bisection Task (Figure 1); correctly predicted that parameters drawn from behavior on a single-duration estimation task would produce geometric bisection in Temporal Bisection Task (Figure 2A); quantitatively predicted shifts in the bisection point produced by varying reward ratios (Figure 2B); and explained why subjects display psychometric reversals for probe durations longer than the “long” reference duration (Figure 2C). Taken together, the framework’s success at characterizing the subjects’ behavior suggests that our rats were not performing the Temporal Bisection Task as traditionally assumed (namely, basing their responses on a comparison of perceptual similarity of a probe to the long versus short references), but were instead producing behavior strongly influenced by temporally-evolving changes in the incentive value of different responses, as caused by temporal discounting of distal rewards.

With the goal of determining a bisection point driven largely by perceptual factors, we therefore sought modify the classic Temporal Bisection Task in a manner that would eliminate temporally-discounted reward value as a driver of response behavior.

We used two variants of the Temporal Bisection Task. The first, which we dubbed the “Immediate Reward” version, was essentially identical to classic versions of the task [21], with rewards baited immediately following the termination of the stimulus (Figure 3A top). In the second version, which we dubbed the “Delayed Reward” version, we simply baited rewards in the appropriate port following a fixed delay from the start of the trial, regardless of whether it was a “short” or “long” trial (Figure 3A bottom; for T_short_=1s T_long_=9s, rewards were baited at T_rew_=10s and for T_short_=100ms T_long_=900ms, rewards were baited at T_rew_=1s). For direct comparison to the classic Temporal Bisection Task, in both variants we measured the bisection point by evaluating performance on intermediate duration probe trials (10% of trials, for which reward was randomly baited on either side port, see Methods). We could not use the same method for estimating the bisection point described above for Figure 1B because in the delayed reward version the stimulus start time was not constant. We trained 12 rats on both variants of the task, and used within-subject comparisons across the two variants.

**Figure 3.**
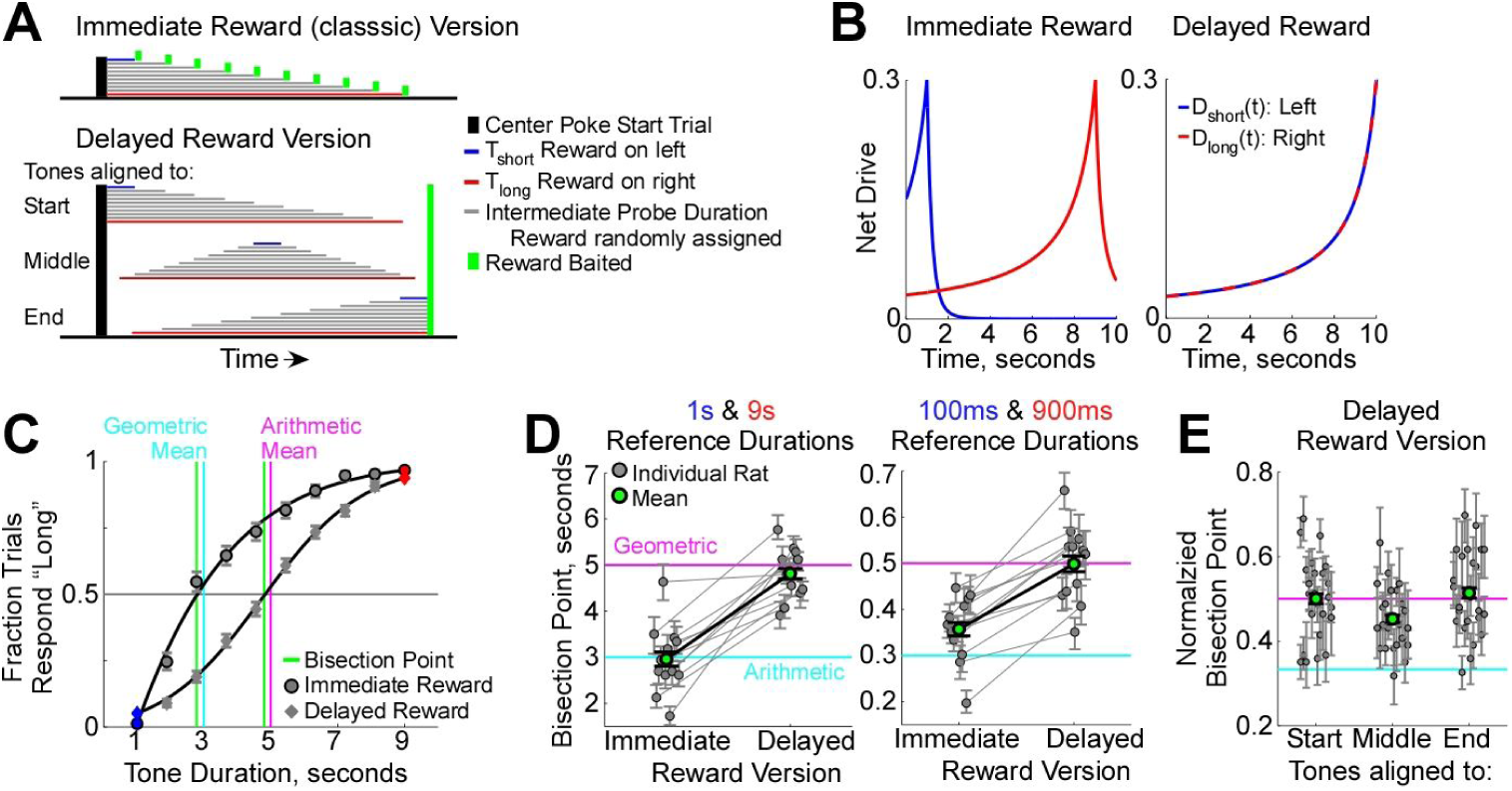
Controlling for temporally discounted reward value shifts rats from geometric bisection to arithmetic bisection. **A)** Top: Immediate Reward (classic) Version of the Temporal Bisection Task. Rewards are baited immediately following the termination of the stimulus. Bottom: Delayed Reward Version of the Temporal Bisection Task. Rewards are baited following a fixed delay from the start of the trial. Stimuli are aligned to 1 of 3 times in the delay period: start, middle, or end. Reference durations T_short_ (blue) and T_long_ (red) account for 90% of trials and are consistently paired with a reward baited in the appropriate nose post. Intermediate length probe durations (gray) account for the remaining 10% of trials and rewards are randomly baited in either nose port. **B)** Hypothetical net drive functions D_short_(t) and D_long_(t) in both versions of the task. In the Delayed Reward Version both rewards are baited at the same delay leading to equivalent net drive functions that perfectly superimpose. C) Psychometric functions from combined data of 12 rats performing both versions of the Temporal Bisection Task outlined in **A** with reference durations T_short_= 1s and T_long_= 9s. **D)** Individual and mean bisection points for rats on both versions of the task with both 1 and 9s, or 100 and 900ms reference duration pairs. Gray lines connect data from individual rats. **E)** Normalized bisection points on the Delayed Reward Version of the task for both reference duration pairs for each of the 3 tone alignments analyzed separately. Error bars represent 95% confidence intervals for individual rats and S.E.M. for combined data. For both **D** and **E**, green data point and error bar shows the mean and standard error across subjects.

As previously discussed, the Immediate Reward version of the task leads to a large temporal asymmetry between the net drive functions D_short_(t) and D_long_(t) associated with producing “short” and “long” responses (Figure 3B left). Consistent with previous findings, for both trained duration pairs tested, which were T_short_ = 1s : T_long_ = 9s and T_short_ = 100ms : T_long_ = 900ms, the bisection point (2.96s +/-0.15s and 357ms +/-14ms respectively, mean +/- standard error cross subjects, Figure 3C,D) was much closer to the geometric mean (3s and 300ms respectively) than the arithmetic mean (5s and 500ms respectively). In contrast, in the Delayed Reward version of the task, there is an equivalent value for T_rew_ in equation 2 for both D_short_(t) and D_long_(t). The net drive functions cancel in equation 1, thus eliminating discounted reward value as a drive to produce one response over the other (Figure 3B right). To perform above chance, subjects must therefore rely on their estimate of the stimulus duration alone. To ensure that the rats were basing their performance on the duration of the tone rather than any other period within each trial, we varied when the stimulus could occur within the delay period. Stimuli could either begin at the start of the trial (immediately following the rat’s center poke marking t=0), be centered in the delay period, or terminate at the end of the delay period (Figure 3A bottom). To control for any effect that moving between side ports during the delay period may have on the bisection point, for the 100ms vs. 900ms experiment the rats were required to fixate their nose in the center port during the delay period (1s). Despite the delayed reward, rats were able to distinguish the “short” and “long” reference durations equivalently between the two variants of the task (blue and red data points in Figure 3C).

In contrast to the geometric bisection found when rewards were baited immediately following the termination of the stimulus, bisection on the Delayed Reward version of the Temporal Bisection Task was much closer to the arithmetic than geometric mean for both trained duration pairs (4.81s +/-0.11s for 1s vs. 9s where rats were free to move between ports during the delay period, and 499ms +/-17ms for 100ms vs. 900ms where rats were required to fixate in the center port during the entire delay period, Figure 3C,D). All rats shifted their bisection point later in time for the delayed reward version of the task (Figure 3D). When analyzed independently, each of the three stimulus alignment times still yielded bisection closer to the arithmetic than geometric mean (Figure 3E). It should be noted that while the model presented above (Figure 1,2) served as the motivation for how to modify the Temporal Bisection Task to uncover rats’ true bisection point, this finding, that delaying the reward to a fixed time following the start of the trial, shifts rats’ bisection to the arithmetic mean, stands independently of that model. In sum, when controlling for the effect of temporally discounted reward value, rat performance shifts significantly away from the geometric mean, indicating once again that discounting of rewards is a key driver of response behavior on the classic Temporal Bisection Task. Moreover, rat performance becomes consistent with human performance on the task, where bisection is typically found near the arithmetic mean [14] (see [16] for review).

## Discussion

For over four decades, it has been repeatedly demonstrated that non-human subjects bisect at the geometric mean [21,29–35] while for over twenty five years it has been shown that humans tend to bisect near the arithmetic mean [14,25–28] (see [16] for review). This seemingly trivial discrepancy could imply profound differences in how humans and non-humans perceive or process temporal information. Here we have demonstrated, however, that this divergence in performance is most likely the result of each species employing different strategies when performing the task, not differences in the fundamentals of their temporal processing: when temporal discounting of rewards was removed as a differential driver of response behavior, the bisection point for rats shifted to being very close to the arithmetic mean, a result paralleling the results found with humans (Figure 3).

We developed a modeling framework that describes subjects’ behavior in temporal processing tasks. By combining a subject’s temporally-discounted value of reward with their internal noisy estimate of time, our modeling framework offers a novel interpretation of non-human subject’s strategy on temporal tasks. Performance on the Temporal Bisection Task has traditionally been conceived of as driven by the subject responding according to which of the “short” versus “long” trained reference durations they perceive the current stimulus duration is closest to. The bisection point (the duration at which subjects are equally likely to respond “short” or “long”), has therefore commonly been interpreted as the duration subjects find to be perceptually equidistant from the two reference durations (though [65] interpret the bisection point as the duration that is equally conditioned to both “short” and “long” responses). This common interpretation fails to explain a number of findings: Why does behavioral data on the Peak Interval Task, which does not involve duration classification, predict geometric bisection (Figure 2A)?; Why does unbalancing the rewards shift the bisection point (Figure 2B)?; and Why do non-human subjects show psychometric reversals beyond the long reference duration ([32,33,35], Figure 2C)? We propose instead that subjects are not explicitly categorizing durations, but rather producing the behavioral response that has the highest current value, following a function that must combine both their sense of time and temporally-discounted reward. The bisection point should therefore be reinterpreted as the duration when the value of the two response options is equivalent, something which can be predicted from a task involving only one response option and should be shifted by altering reward magnitude.

Work involving a range of species across multiple tasks has demonstrated that reward is one of the leading influences on performance during most behavioral tasks [44,51,66–68], When rewards are terminated altogether, most animal subjects will quickly cease performing any task [69]. When rewards become unbalanced, subjects will favor choosing the option that gives them the larger reward [68,70]. Despite it being such an influential variable, reward value is usually omitted from behavioral models for tasks not explicitly designed to probe it, e.g. discounting tasks or any task that requires the subject to learn about the magnitude of the reward, as researchers go to great lengths ensuring that rewards are both balanced (in multiple choice tasks) and are of a sufficient level to motivate performance without leading to overt satiation. However, temporal tasks, such as those described here, where the delay to rewards may be different, or where response performance is monitored during the course of the entire trial [71], lead to rewards whose incentive values are inherently unbalanced, due to temporal discounting. As a reward becomes more imminent within a trial, the subjective value of that reward should rise. We therefore thought it only logical that a subject’s response behavior should vary accordingly.

By simply delaying the reward until a fixed time following the start of each trial, we were able to eliminate the temporal asymmetries, and thereby remove the ability to use discounted reward value as a strategy to perform the Temporal Bisection Task. In doing so we revealed the true underlying duration categorization behavior of rats. Rats, just like humans, tend to partition durations around the arithmetic mean of two trained reference durations, implying that the neural mechanism by which we perceive and categorize time is likely to be conserved at least amongst mammals.

Despite its ability to account for moment-by-moment response behavior across three disparate temporal processing tasks (see supplemental text and supplemental figures 1-5 for fits to the Delay Discounting and Peak Interval Tasks), our modeling framework is obviously incomplete, as it makes no claim to how subjective time itself is measured [3,5,41,72–74], except that it should be proportional to real time [8], or how durations are learned [1,65,75], except that they should possess scalar noise [36]. Answers to these and many other questions will lead to a more complete understanding of how temporal information is processed in the brain. However, knowing that rats can categorize durations in a similar manner to humans makes findings from non-human subjects all the more relevant in understanding how we perceive the passage of time.

## Methods

### Rat housing and general training

Male Long Evans rats were pair housed in Technoplast cages on a reversed dark/light cycle. Rats had free access to food but had restricted water access limited to 30 minutes per day (starting 30 minutes following the end of training) and what they could earn during training, as approved by Princeton University IACUC. Training duration ranged between 90 and 110 minutes per day and occurred at the same time each day, seven days per week. Behavioral chambers (Island Motion) consisted of a raised grated floor, three conical nose ports on one curved wall, and stereo speakers. Each nose port contained one LED that could be illuminated according to task or training requirements, an IR beam to detect nose pokes, and water tubes for reward delivery (only in the left and right ports). Water was fed via gravity from a reservoir. Reward volume was controlled by the valve open time. Each chamber was housed within a sound attenuation box (Coulbourn). Training was fully automated utilizing the Bcontrol System.

### General Training

Training of all tasks began with the following stages designed to familiarize the rat with the training environment. 1) The first day rats learned that poking into the left port illuminated by an LED produced a drop of water 24µl. On the following day they learn the right port. 2) Rats learned to switch between poking in the left and right port within a session. 3) Rats must make a center poke to initiate the trial and then poke in whichever side port was illuminated.

### The Modified Temporal Bisection Task

Training continued from general training: 1) Rats learned to associate the short duration (0.39, 0.78, 1.57, or 3.17s) with reward in the left port and the long duration (10x the short duration) with reward in the right port. The center LED cued that the trial was ready, and the rat must make a center poke to initiate the trial. An 11.3 kHz tone of either the short or long duration began 50ms following the start of the trial after which rewards were baited. The rat’s response was taken as the first side poke following reward baiting. On any one trial only 1 reward was baited. Incorrect responses elicited 8s of white noise and no water reward. Trials were in blocks or 40 alternating between short duration left reward baiting and long duration right reward baiting. 2) Long and short duration trials come randomly. Once a rat can perform a minimum of 100 trails within a single session with performance on both short and long duration trials at or above 80% they began data collection. Full Task: 3) 150 long duration trials were collected. This was combined with the long duration trials from the last session in stage 2 for model fitting. 4) Repeated stages 1-3 for each of the other duration pairs. Rats progressed through the pairs in random order.

### Immediate (classic) and Delayed Reward versions of the task

Training was the same as the modified Temporal Bisection Task outlined above for stages 1-2. Eight intermediate probe durations were introduced randomly on 10% of the trials, spaced linearly between the trained short and long reference durations. Rewards were baited randomly between the left and right port on probe duration trials. For the immediate reward version 60 trials of each of the 8 probe durations were collected from each rat. For the delayed reward version the tones were aligned to either the start, middle or end of the delay period and 40 trials of each of the 24 probe trial types (8 durations x 3 alignments) were collected from each rat. For the 100ms vs 900ms discrimination on the delayed reward version rats were required to keep their nose in the center port for 1s during which time the tone played (the center light remains illuminated until the rat is allowed to leave). Violation trials in which the rat leaves the center port for more than 100ms are excluded from analysis. A response in the nose port with the baited reward earns the rat 24µL of water while a response in the opposite port earns it no water and 8s of white noise. Any responses elicited before the reward is baited are ignored.

### DDM fitting

DDM simulations were run in MatLab. Parameter fitting was performed following a gradient ascent using MatLab’s fmincon function for each experiment independently to maximize the log-likelihood of producing all the poke rasters within the experiment. 25 randomly seeded gradient ascents were run per experiment to fit the model’s free parameters R, k, s, h, f, see equations 1 and 2). For this first run of the fitting the values of T_rew_ in D_short_(t) and D_long_(t) were set to the actual reward times in the experiment T_short_ and T_long_, respectively. The best fit parameter values were then held constant and the best fit values for T_rew_ were found for each trial. The gradient ascent then iterated between fitting the model parameters and T_rew_ until no further improvement in the log-likelihood was achieved.

### Model Framework and the Center Nose Port

Each of the tasks presented here began when the rat self-initiated a nose poke into the center port. The nose in event was what was counted as t=0 for each of the trials, not the nose out event. We modeled the center poking just as we modeled the left and right poking behavior. At the start of each trial the model was assumed to be in the center nose port (as it must be to start a trial). The duration of that poke was taken randomly from the center poke events produced by the rat during that particular experiment. Following the completion of the center poke the DDM was set to 0 (regardless of the hysteresis parameter) and started running. No further center poke events were allowed by the model during a trial and any rare center pokes produced by the rat during the trial were ignored when calculating the log-likelihood.

### Calculating bisection from the Modified Bisection Task

Poke rasters were produced from the poke in and poke out times with 20ms resolution. The inter-poke intervals were interpolated by the previous and subsequent pokes meeting at the arithmetic mean of the previous pokes out time and the subsequent pokes in time. IPIs that occurred at the end of a trial, i.e. they were not flanked by two pokes, were left as NaN, as were the center pokes initiating the trial. Left pokes were converted to 0 and right pokes were converted to 1. The mean was then taken across trials (excluding NaNs) to produce the psychometric function of “long” reports as a function of time.

### Calculating bisection from the Classic Bisection Task

A plot of the fraction of “long” responses as a function of stimulus duration t was fit with a four parameter logistic function

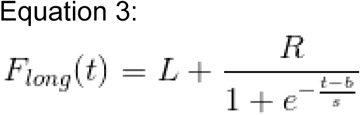

from which the bisection point (the duration that produced 50% “long” responses, F(t_bisection_) =0.5) was extracted.

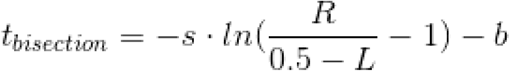

L represents the lower bound, R the range between the lower and upper bound, b the bias, and s the slope of the function. 95% confidence on bisection was estimated by bootstrapping 10,000 times: sampled trial response pairs with replacement and then refit the logistic function and extracted the bisection point.

## Supplemental Text

The model we present in the main text was conceived as a general framework translating a subject’s sense of time and reward into task relevant actions. For time intervals spanning seconds to minutes, two largely separate bodies of research focus on delay discounting [44–54] and time estimation [1,8,14,22,36,38–43]. Delay discounting experiments assay subjects’ choices given rewards delivered at different delays. Non-human subjects learn about these delays from experience over trials– they must make time estimates of the delays involved. Conversely, in time estimation experiments, subjects must estimate delays, after which rewards are delivered. According to the delay discounting literature, the value of these rewards will be strongly affected by the length of the delays.

Here we take a step towards merging these two lines of research (see [76,77], for other efforts in bridging these two literatures) by presenting a phenomenological and behavioral modeling framework that readily combines the core concepts of both literatures. The model incorporates reward value, discounting of future rewards, and noise in time estimation. Decisions about action are further assumed to have a stochastic component, which allows the fitting of moment-by-moment and trial-by-trial response behavior.

We trained rats in three classic yet distinct temporal processing tasks: Delay Discounting [44], Peak Interval [62], and Temporal Bisection [21] (see main text). All three were well fit by the same modeling framework, with parameter values indicating that noise in time estimation and discounting are important in each. Since the framework can be thought of as a behavioral strategy, we propose that despite substantial differences in the definition of these tasks, a similar overall strategy may be used to perform them.

In order to capture the full dynamics of a subject’s real-time response behavior, a model must be capable of replicating both the timing of response initiations and the duration of each response. In all of the experiments we performed, we found that the duration of time for which responses were elicited (in our case, duration of time over which rats’ noses remained in the nose ports, once they had poked into them) did not depend strongly on time within a trial or any other experimental parameters (Supp Figure 1A). In contrast, the rate at which responses were initiated did depend very strongly on experimental parameters (Supp Figure 1B). Consequently, we focused our modeling efforts on the intervals between completing an actuator operation (withdrawal from a nose port) and initiating another response (start of next nose poke). The duration of nose pokes themselves were modeled by simply randomly drawing durations from the subject’s own data, disregarding information about the trial or time elapsed during the trial.

**Supp Figure 1.**
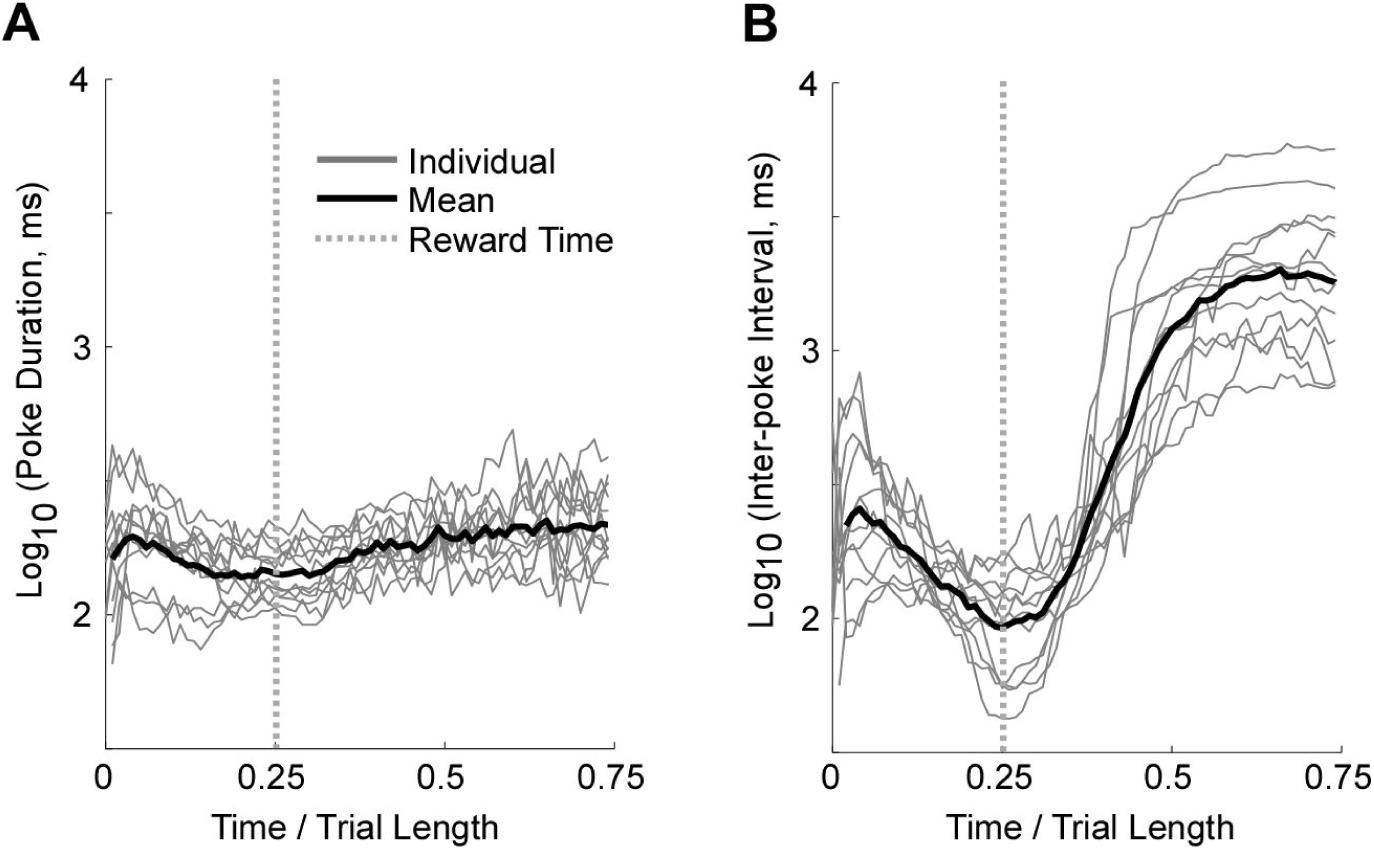
Inter-poke Interval not poke duration varies as a function of time within trials. **A)** Average poke duration as a function of time during the Peak Interval task on reward omitted trials. **B)** Average inter-poke interval (IPI) as a function of time during the Peak Interval task on reward omitted trials.

### The Delay Discounting Task

We trained 12 rats in a modified version of the Delay Discounting Task [44,54]. Each trial began with the onset of a light-emitting diode (LED) located in the center port (Figure 1A), indicating that a new trial was ready. Rats were trained to respond to this LED by poking their nose into the center port; this event marked time t=0 seconds (Supp Figure 2A). Rats were then free to poke into any port they wished. Two types of pokes could terminate trials: (1) after a time T_short_, a poke into the left nose port (which we called the “soon” port) produced a small water reward, followed by a delay and trial termination; (2) after a time T_long_, a poke into the right nose port (which we called the “late” port) produced a large water reward, followed by a delay and trial termination. The rats were thus making a choice between receiving a small reward quickly at the “soon” port, or receiving a larger but more delayed reward at the “late” port. Post-reward delays were chosen so that the total trial length was the same for either choice (60 seconds). The small reward was always a single 24 µl drop of water, and T_short_ was always kept fixed at 2 seconds. T_long_ was adjusted after each trial until the rats chose the “soon” and the “late” ports with equal probability (see Methods, [54,78]).

Each rat was a subject in two separate experiments: in one the large reward was 2 drops of water, and in the other the large reward was 4 drops of water. Supp Figure 2B shows the stable values of T_long_ found for all 12 rats in both experiments. As has been commonly found in delay discounting experiments, the larger the reward, the longer subjects are willing to wait for it [44,53,54,61]. If X drops of water after time T_long_ are valued equally to 1 drop of water after time T_short_, then the delay discounted value of a drop of water is said to have been reduced by a factor 1/X (the discounted value, DV, equation 4). We plot the relative delay discounted value as a function of T_long_ in Supp Figure 2C. Again as commonly found in delay discounting experiments, the data are well fit by a hyperbolic function (r = 0.95) [44,47,53,54,61]

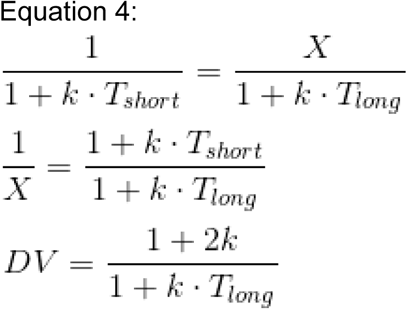

Equation 4 is written such that the relative discounted value DV for X drops at T_long_ seconds equals 1 and is equivalent to 1 drop at T_short_ = 2 seconds. The parameter k, which we fit separately for each rat, is known as the discounting constant.

**Supp Figure 2.**
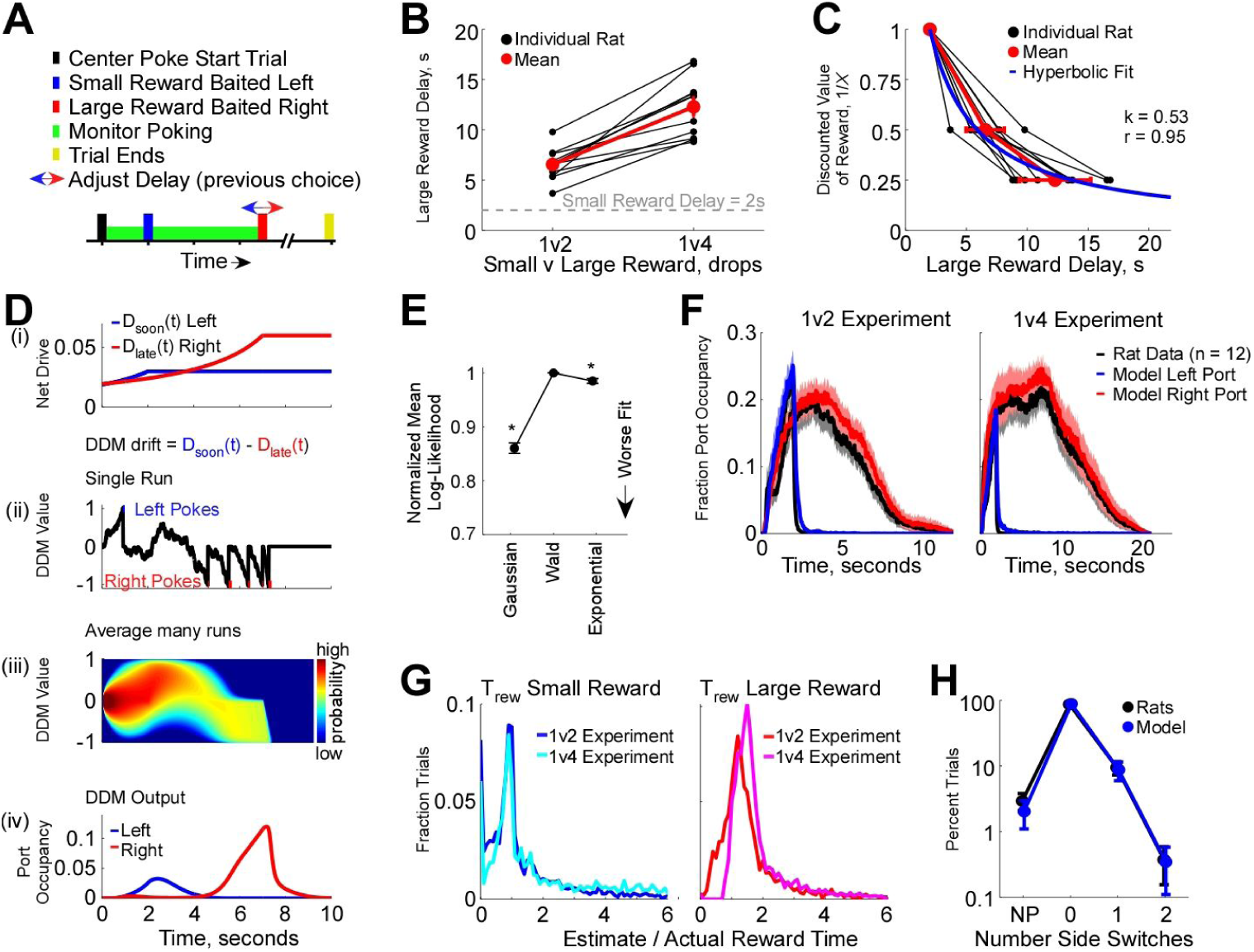
The Delay Discounting Task. **A)** Task structure. **B)** Large reward delays that lead to equal selection of the large (2 or 4 drops) and small (1 drop) rewards. **C)** Large reward delay (from **B**) plotted against the discounted value of the large reward. Hyperbolic fit, equation 4, uses the median discounting constant calculated from individual rat fits. **D) i:** Example hyperbolic net drive functions for the left and right ports. **ii:** Single run of the DDM with the left and right nose pokes produced. **iii:** Fokker-Planck solution for the DDM showing full probability distribution of DDM values. **iv:** Response probability taken from threshold crossing in **iii. E)** Log-Likelihood of best fit Gaussian, Wald, and exponential distributions fit to IPI histograms taken from discrete 2s time periods across trials. Fits computed separately for each time window for each rat. * for p<0.05 **F)** Mean left (“soon”) and right (“late”) response probability for the 1v2 and 1v4 drop experiments averaged across all rats (black) and model fits (color). **G)** Histogram of T_rew_ parameter used to fit individual trials combined across all rats separated by experimental condition. T_rew_ is normalized by the actual reward time in the experiment. **H)** Percent of trials on which various number of side switches were made. NP: no pokes elicited before both rewards are baited. Note the logarithmic scale on the vertical axes. Error bars S.E.M.

As described in equation 4 and in the delay discounting literature, the larger the delay between the current time and the time of a future reward, the less that reward is valued. Logically then, as a reward approaches in time within a trial its subjective value should rise following the same hyperbolic function. We thus rephrase equation 4 as the statement that at time t, the net drive D(t) for X drops of water that will become available at time T_rew_, and will then remain available until harvested, is equal to

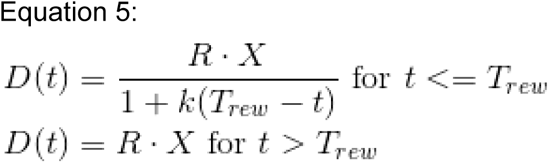

where R is a scaling constant. For the period t <= T_rew_ equation 5 is identical to equation 2 presented in the main text. However, because both rewards remain available in this task until one is earned, the net drive does not decay following T_rew_ but rather remains constant.

Based on the time estimation literature [1,8,14,22,36,38–43] we know the subject’s estimate of T_short_ and T_long_ will be subject to noise. Behavioral models typically account for this noise by describing the estimate of a duration as a distribution from which a unique value is pulled on each trial [14,39], Translating this back to the subject, the rat should therefore have a discrete and unique value for T_short_ and T_long_ on each trial and behave accordingly. For example, on trial 10 the subject’s estimate of T_short_ may be 2.5s and for T_long_ may be 6s while on trial 11 they may be 1.7s and 5.1s respectively. Here we are fitting our model to the moment-by-moment trial-by-trial poking behavior (Supp Figure 3) rather than the summary choice statistics (did he earn the small or large reward) or the average response profile (response probability Supp Figure 2F or mean response rate). In order to determine the model parameters that maximize the likelihood of replicating the exact nose-poking pattern produced by the rats for each trial, we must know the value that was pulled for T_short_ and T_long_ on each trial. It is not enough to know that the estimate of T_short_ follows a certain distribution with a specific mean and standard deviation, since the trials the model performs would not correspond to the trials the subject performs. Following the example above, the model by chance may pull a value of 4s for T_short_ on trial 10 causing it to produce a different response pattern from the subject, leading to a poor fit, but not because the model is necessarily bad but because the stocasticity of the model and the rat are not in sync. Therefore, since the subject likely draws a discrete value from their noisy estimate for T_short_ and T_long_ on each trial, we found those values that maximize the log-likelihood of the model replicating the subject’s response behavior (see Methods). In doing so we not only avoid making an assumption as to what type of distribution should describe the subject’s estimate (Gaussian, log-normal, gamma…) but also can directly assess that distribution by simply plotting the histogram of the best fit values for T_short_ and T_long_ across trials (Supp Figure 2G). By maximizing the likelihood of the model replicating the precise poking behavior produced by the subject in a moment-by-moment and trial-by-trial manner (at 20ms resolution for both the left and right nose ports, see Methods), rather than simply their final “choice”, we are attempting to fit 100 data points per second per trial that the subject performs. Finding the best fit values for T_short_ and T_long_ on each trial therefore does not lead to over-fitting as the number of fit parameters per experiment is always orders of magnitude below the number of data points we are attempting to fit. On average the delay between the start of the trial and delivery of a reward is 5.85s. There are therefore ~290 data points per fit parameter for the Delay Discounting Task.

Here we used the trial-by-trial estimate of T_short_ as T_rew_ and set X=1 in equation 5, and used the resulting expression as the net drive function D_soon_(t) for poking into the “soon” port in equation 1. Similarly, we used the trial-by-trial estimate of T_long_ as T_rew_ and set X=2 or X=4 in equation 5, according to the size of the large reward in each experiment, and used the resulting expression as the net drive function D_late_(t) for poking into the “late” port. Supp Figure 2D illustrates the behavior of the resulting model. In this hypothetical example trial the model produces a single left poke prior to either reward being baited and therefore does not terminate the trial. Later in the trial the net drive for the “late” port dominates driving the model to switch sides and produce right pokes. Supporting our use of a DDM, histograms of the inter-poke intervals taken at fixed times across trials are better fit by a Wald distribution than either Gaussian or Exponential distributions (Supp Figure 2E).

Poke rasters for each rat performing 300 trials of the 1 vs. 2 drop experiment are shown in Supp Figure 3A and the 1 vs. 4 drop experiment in Supp Figure 3B alongside the output of the model fit to this data (1v2 experiment mean r^2^ = 0.93, 1v4 experiment mean r^2^ = 0.95) with all pokes following the earning of a reward omitted (not used in the fitting). The fraction of trials in which a port is occupied at any given time is obtained by averaging the left and right pokes independently, and serves as a convenient means for comparing the rat and model’s response behavior. Supp Figure 2F shows the response probability averaged across all 12 rats for each of the two experiments along with the average model fits. Here k in the model (Equation 5) is not a fit parameter but rather calculated from each rat’s choice behavior (Supp Figure 2B,C).

Traditionally, time estimation errors have not been considered in Delay Discounting Tasks. Here, however, T_short_ and T_long_ were best fit by broad distributions (Supp Figure 2G), suggesting that such errors play a significant role in determining behavior. In addition, the model fits detailed characteristics of the poking, including the frequency of switching sides. At the start of most trials both the rats and the model choose either the “soon” or “late” port, making repeated nose pokes there until a reward is earned (88.6% of trials for the model, 87.1% for the rats, Supp Figure 2H). However, on a small subset of trials (2.0% for the model, 2.9% for the rats) both the rats and the model produce no pokes at all (NP) before the late reward is baited. On a larger percent of trials (9.1% for the model, 9.5% for the rats), both the rats and the model make a single side switch before a reward is earned. Both the rats and model occasionally make 2 side switches (0.2% for the model, 0.4% for the rats), however, neither switch sides more than twice in the same trial (Supp Figure 2H).

To our knowledge, this is the first attempt to model moment-by-moment trial-by-trial response behavior in a delay discounting task.

**Supp Figure 3.**
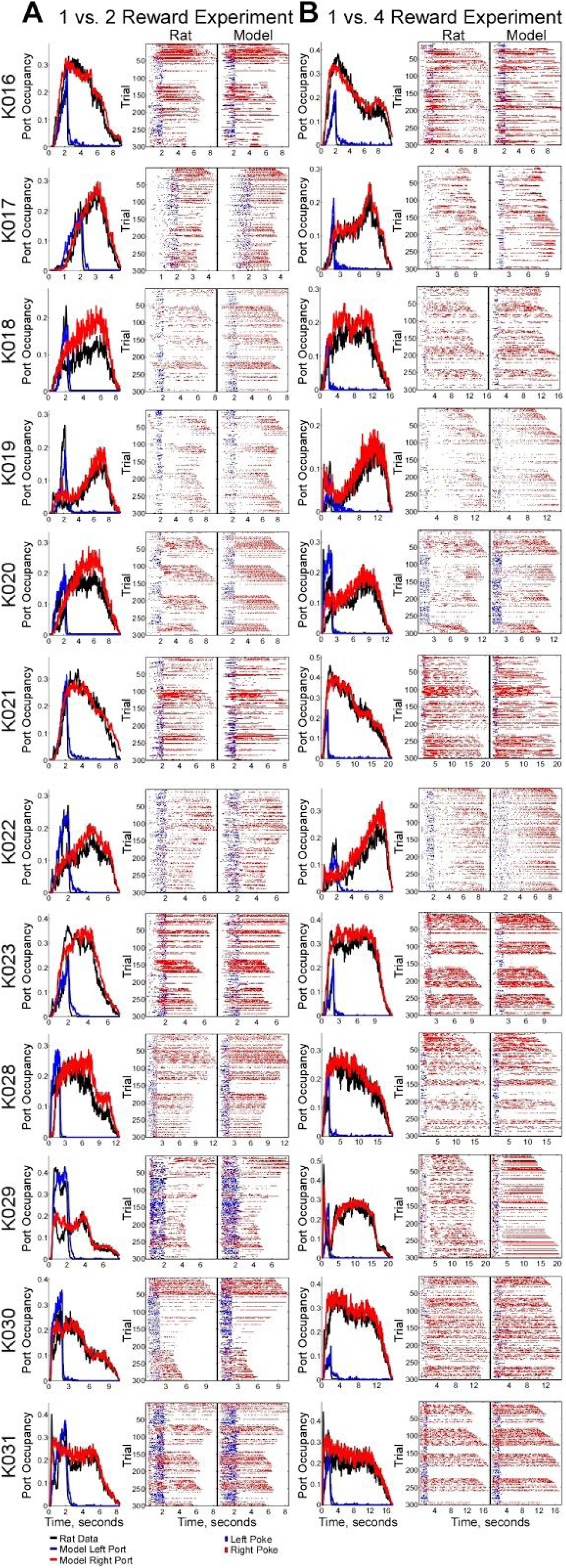
Individual rat performance and model fits for the Delay Discounting Task. **A)** Left column: Response probability as a function of time for the rat (black) and model (color) on the 1 vs 2 drop experiment. Center column: Individual rat poke rasters. Right column: Poke rasters produced by the model fit to the individual rat’s data. **B)** Same as **A** for the 1 vs 4 drop experiment.

### The Peak Interval Task

To test whether the form of drive that we used to fit Delay Discounting data (equation 5) could also be used in a different, unrelated task, we trained rats in an often-used temporal processing task, the Peak Interval Task [38,39,62,79–82]. As with the Delay Discounting Task, each trial in our Peak Interval Task began with the onset of an LED located in a center port, indicating to the animal that it could start a new trial (Supp Figure 4A). Trained rats responded to this by poking their noses into the center port, which turned the LED off; this event marked time t=0 seconds. Rats were then free to poke into any port they wished. On rewarded trials (a randomly chosen 85% of trials), the first poke after a time T_rew_ into the left nose port resulted in a 24µl drop of water. After 4x T_rew_ seconds had elapsed, a new trial began, indicated by turning the center LED on again.

**Supp Figure 4.**
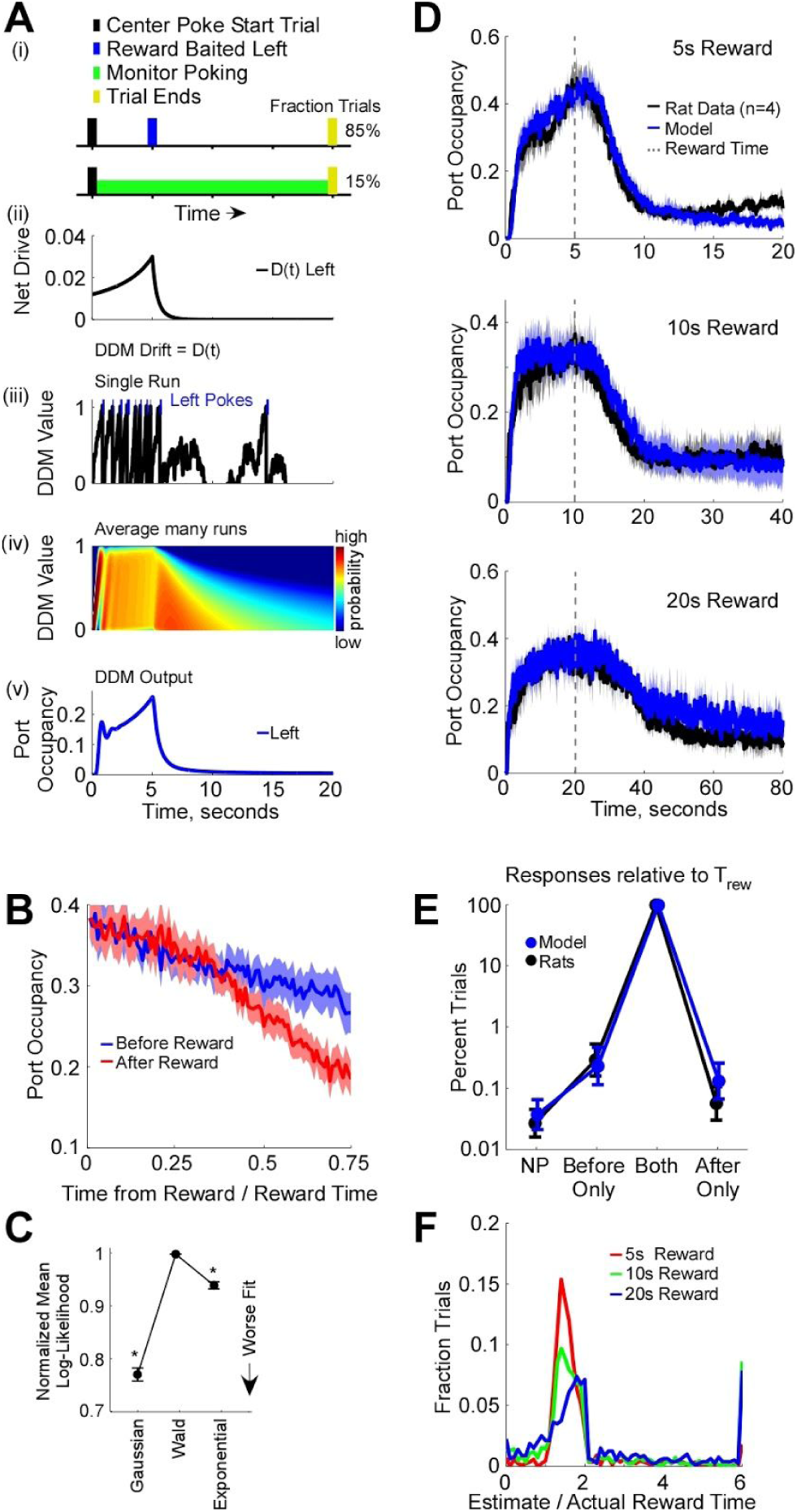
The Peak Interval Task. **A) i:** Task structure. **ii:** Hyperbolic net drive function for the left nose port following equation 7 for the pre-reward and post-reward period. **iii:** Single run of the DDM with the left nose pokes produced. **iv:** Fokker-Planck solution for the DDM showing full probability distribution of DDM values. **v:** Response probability taken from threshold crossing in **iv. B)** Mean response probability as a function of time from the reward either before (blue) or after (red) the reward time (on unrewarded trials). **C)** Log-Likelihood of best fit Gaussian, Wald, and exponential distributions fit to IPI histograms taken from discrete 2s time periods across trials. Fits computed separately for each time window for each rat. * for p<0.05 **D)** Mean rat (black) and model (blue) response probability as a function of time for the 3 reward conditions tested (5s, 10s, and 20s) on non-rewarded trials. **E)** Analyzing the period 0<t<2x T_rew_ on unrewarded trials, percent of trials that contain no pokes (NP), pokes only before T_rew_ (Before Only), pokes before and after T_rew_ (Both), and pokes only after T_rew_ (After Only). **F)** Histogram of the T_rew_ parameter used to fit individual trials combined across rats separated by experimental condition. T_rew_ is normalized by the actual reward time in the experiment. Error bars S.E.M.

As is typically observed in Peak Interval Tasks [38,39,62,79–82] the response rate (here, nose poking), increased as T_rew_ approached (Supp Figure 4B). On the main trials of interest, which are the randomly-chosen 15% of trials in which the reward was omitted, the response rate peaked around T_rew_, and then gradually decayed afterward (Supp Figure 4B,D). The peak near time t≅T_rew_ indicates that subjects can estimate the time in the trial relative to T_rew_, and must therefore have some internal memory of T_rew_ [62]. The width of the peak around T_rew_ is usually taken as indicative of the subject’s error in their estimate of T_rew_ [38,80,82]. As previously observed, both in Peak Interval and many other timing tasks, this width scales approximately linearly with T_rew_ (Supp Figure 4D,F). This is referred to as scalar timing [63,80]. Following the Peak Interval literature, we focused our analysis on unrewarded trials. Note that unlike the Delay Discounting Task-- in which once reward becomes available, it remains available until harvested or the trial ends-- after time T_rew_ the absence of a reward indicates that this may be an unrewarded trial, and the reward will never become available. Below, this difference with respect to Delay Discounting will be reflected in a modification to the model’s net drive function for times t>T_rew_.

As a reward approaches in time within a trial its subjective value should rise following the same hyperbolic discounting function discussed above for the Delay Discounting Task. We therefore used the same modeling framework to fit the Peak Interval Task data. We first used equation 5 for t<=T_rew_ and fit the model to the individual poke rasters within the interval 0<t<=T_rew_. On average, the model using equation 5 for t<=T_rew_ is able to fit the rat behavior well in this time period (Supp Figure 4D, t<=T_rew_). Then, to find an appropriate net drive function for t>T_rew_ on unrewarded trials, we examined the rate of poking after T_rew_. Supp Figure 4B shows a comparison of response probability before vs. after T_rew_ on unrewarded trials. Overall, the response rate falls following T_rew_, and does so at a greater rate than it rises before T_rew_ (p<0.01).

Two possible empirical modifications to equation 5 that would produce the asymmetry of Supp Figure 4B are shown below. First, in equation 6 we consider the possibility of a different discounting constant k’ for t>T_rew_:

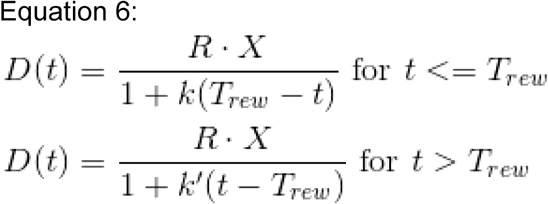

Second, in equation 7 we use the same discounting constant k for t<=T_rew_ and for t>T_rew_, but we add an exponential term in the denominator for t>T_rew_.

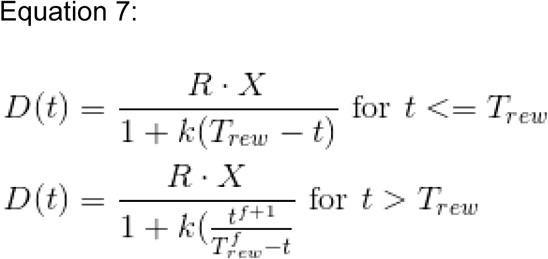

The greater the value of f the more rapidly the net drive function falls off following T_rew_. Note that equations 6 and 7 have the same number of free parameters, and X in these experiments is fixed at 1.

We found that, on average, equation 7 fit the rat response behavior better than equation 6 (p<0.01). Equation 7 also fit the data better than a Gaussian function (p<0.01), which would be symmetric around T_rew_, and better than the case where net-drive immediately falls to zero following T_rew_ (p<0.01).

**Supp Figure 5.**
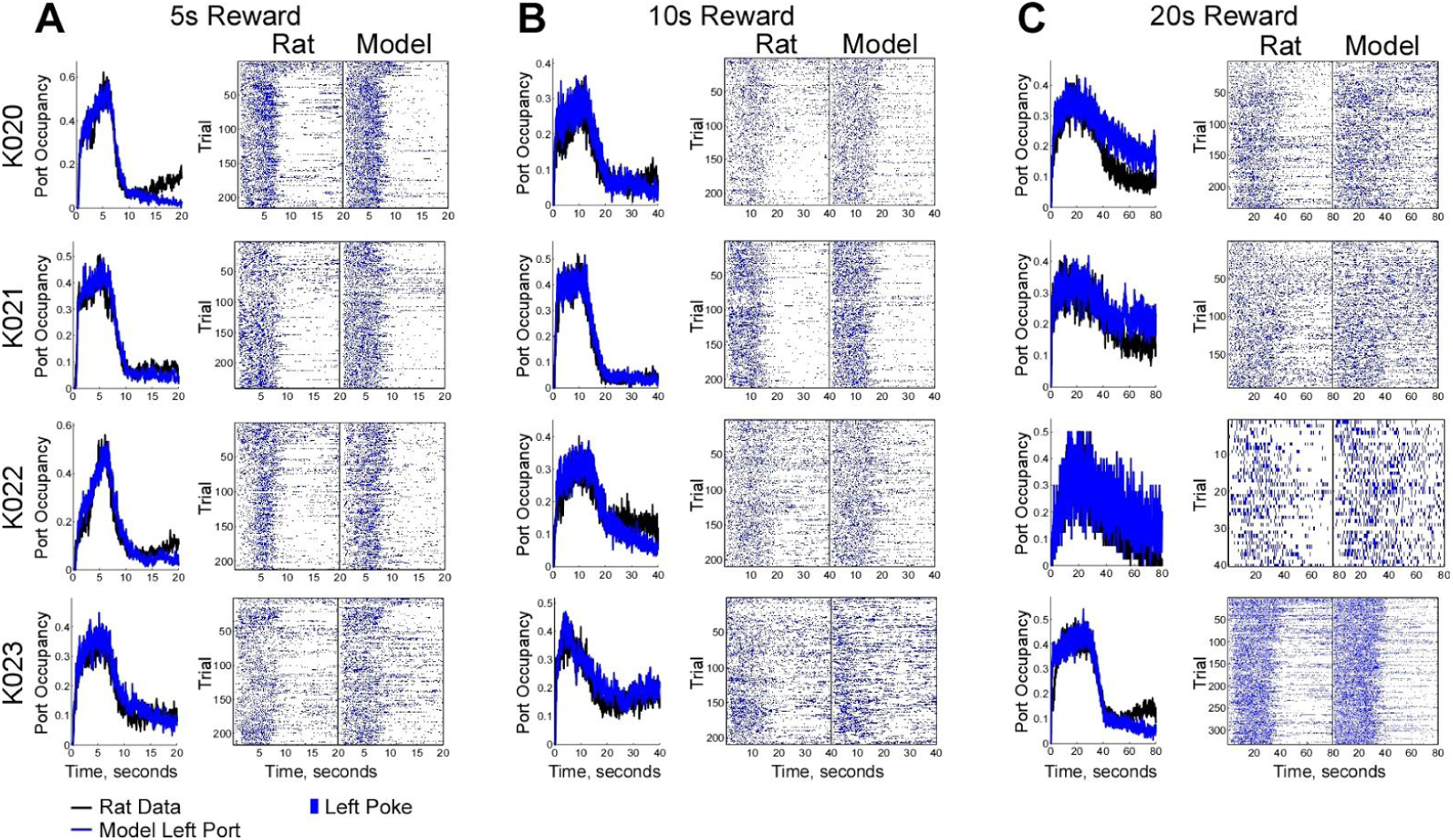
Individual rat performance and model fits for the Peak Interval Task. **A)** Left column: Response probability as a function of time for the rat (black) and model (blue) on the 5s experiment. Center column: Individual rat poke rasters. Right column: Poke rasters produced by the model fit to the individual rat’s data. **B)** Same as **A** for the **1**0s experiment. **C)** Same as **A** for the 20s experiment.

Supp Figure 4A demonstrates how the model framework would behave given a net drive function following equation 7. Supp Figures 4D,E and Supp Figure 5 show that it can fit both mean behavior and moment-by-moment trial-by-trial response behavior well, respectively. As we have shown for the Delay Discounting and Temporal Bisection Tasks, again supporting our use of a DDM, histograms of the inter-poke intervals taken at fixed times during repeated identical trials are better fit by a Wald distribution than either Gaussian or Exponential distributions (Supp Figure 4C). In Supp Figure 4F we plot the histograms of the trial-by-trial estimates of T_rew_ (see explanation in the Delay Discounting Task section for why T_rew_ is fit in this manner), normalized by the actual reward time. The distributions resemble Gaussians but also possess a very long tail in the positive direction. In addition, the distributions are all shifted to have medians greater than the time when the reward is actually baited (medians relative to T_rew_ are 1.54, 1.62, and 1.78 for the T_rew_=5, 10, and 20s experiments respectively). This makes sense as the rat must poke after the reward is baited to earn it. Finally, the distributions appear very similar for the three experiments, consistent with the scalar timing literature [63,80].

In this task, rats will produce nose pokes into the rewarded port on most individual trials both before and after T_rew_. However, on a small fraction of trials rats omit responses by either not poking around T_rew_ (period 0<t<2T_rew_, “NP”), poking before T_rew_ but not after T_rew_ (no pokes within T_rew_<t<2T_rew_, “Before Only”), or poking only after T_rew_ (“After Only”). Once again, the model is able to capture these anomalous behaviors (Supp Figure 4E) not as a failure of attention or engagement in the task but rather as a consequence of the stochastic decision making process and the noise in the estimate of T_rew_.

**Supp Figure 6.**
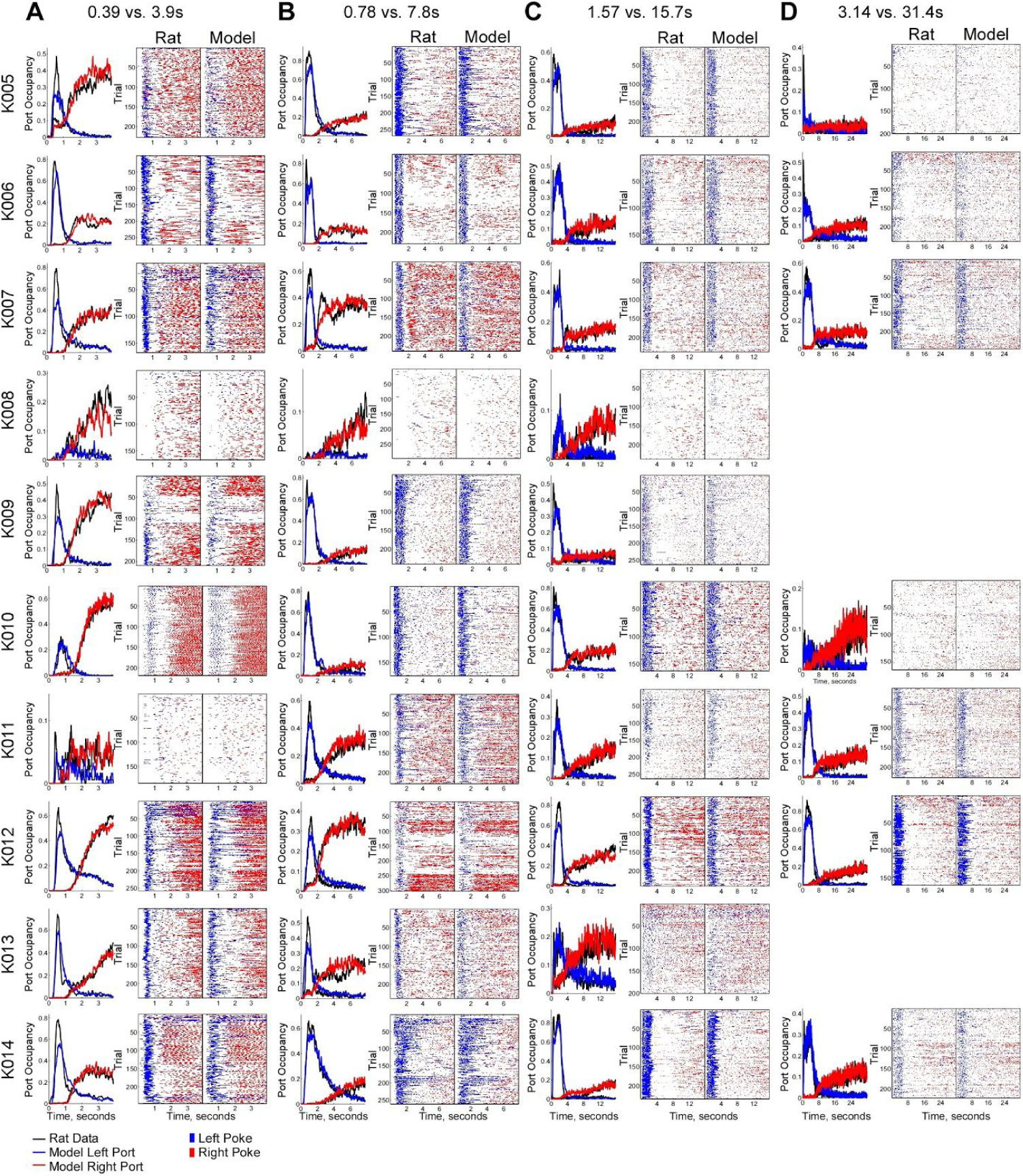
Individual rat performance and model fits for the Temporal Bisection Task. **A)**Left column: Response probability as a function of time for the rat (black) and model (color) for duration Pair 1 (T_short_ = 0.39, T_long_ = 3.9s). Center column: Individual rat poke rasters. Right column: Poke rasters produced by the model fit to the individual rat’s data. **B)** Same as **A** for duration pair 2 (T_short_ = 0 78, T_long_ = 7.8s). C) Same as **A** for duration pair 3 (T_short_ = 1.57, T_long_ = 15.7s). **D)** Same as **A** for duration 4 (T_S_ho,t = 3.14, T_long_ = 31.4s). Data was not collected from 3 rats for technical reasons.

**Supp Figure 7.**
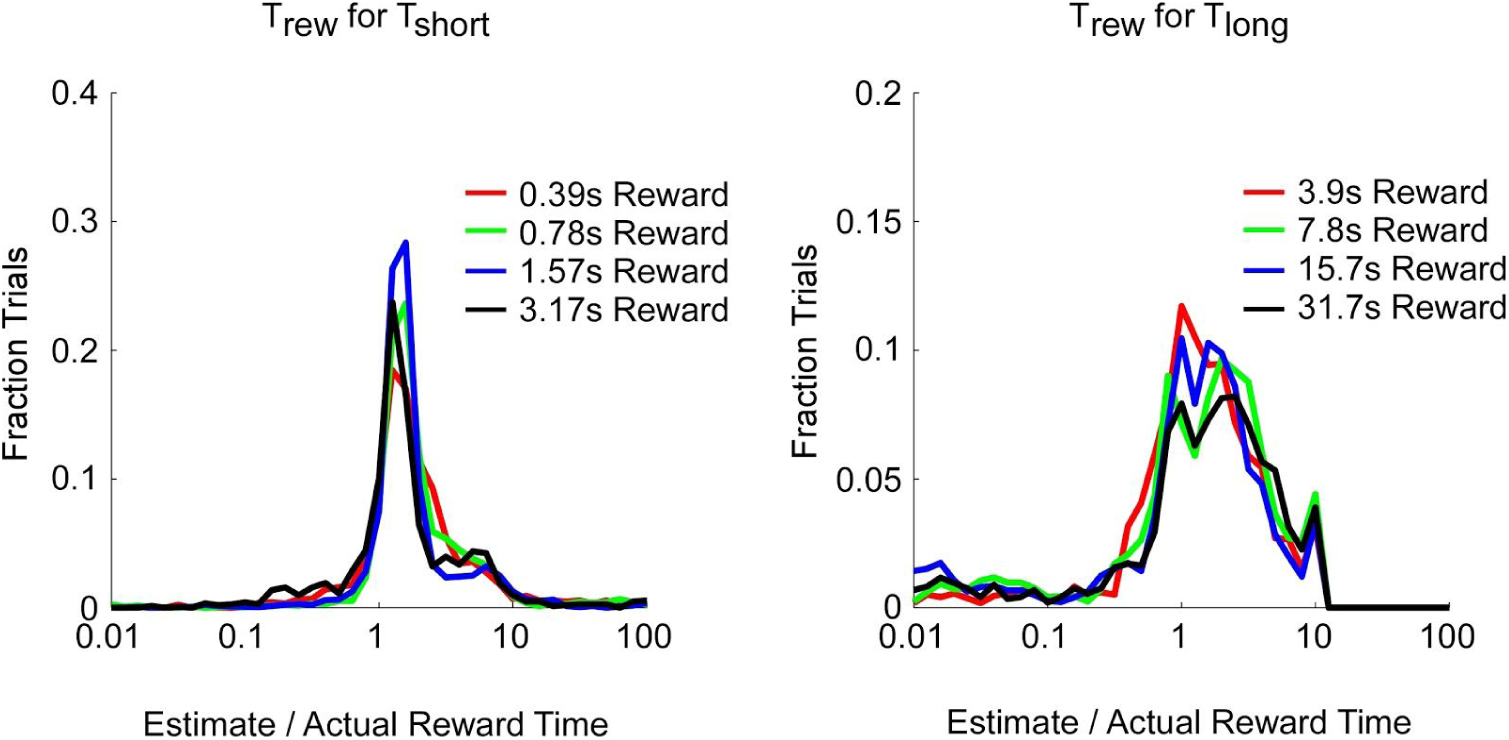
Estimates of reward time in the Temporal Bisection Task follow scalar distributions. **A)** Histogram of T_rew_ fit for I T_short_ on individual trials combined across all rats separated by duration pair. **B)** Same as **A** but for T_rew_ fit for T_long_. T_short_ and T_long_ are normalized by the actual reward time in each experiment. Note the log scale on the horizontal axis.

## Supplemental Methods

### The Delay Discounting Task

Training continued from general training (see Main Methods): 1) Alternating left and right trials in blocks of 10. The reward was baited 2s after the start of each trial (1 drop left, 2 drops right). An LED illuminated the side port with the baited reward. Trial length was kept fixed at 60s. Full Task: 2) Trials come in blocks of 5: only left baited, only right baited, then 3 trials where both were baited following their respective delays. Side LEDs turned on following start of the trial for the side(s) that were baited during that trial. Rats were free to switch sides until a reward was earned. Large reward delay (LRD) began at 2s (equivalent to the small reward delay) and was then adjusted based on the rat’s choice on the previous trial (LRD x 1.01 following a large reward choice, LRD / 1.01 following a small reward choice). 500 free choice trials were then collected and the model was fit to the last 300, which gave the large reward delay enough time to reach a steady state. 3-4) Same as 1 and 2 but with 4 drops baited on the right.

### The Peak Interval Task

In this task the center LED was illuminated when trial was ready, a center poke turned the center LED off and turned on the left LED marking time t=0s. Training continued from general training: 1) 200 trials performed with trial length 10s and reward baited immediately. 2) 300 trials performed with trial length 20s and rewards baited after 5s. Full Task: 3) 15% of trials introduced with no reward baited. Data collected from 200 non-rewarded trials. 4-5) same as 2-3 but with trial length 40s and rewards baited at 10s. 6-7) Same as 2-3 but with trial length 80s and rewards baited at 20s.

### Fitting k for Delay Discounting data

k from equation 4 in Supp Figure 2C was fit for individual rats with a gradient ascent. The median value of k across rats was used as the fit for the average rat data. For the Delay Discounting experiments k was not fit as a free parameter in equation 5 in the model but kept fixed as the value calculated from their behavior (Supp Figure 2C) according to equation 4.

## Acknowledgements

We thank J. Teran and L.K. Osorio for all rat handling, J. Erlich, B. Brunton, J. Jun, K. Miller, and members of the Brody Lab for comments on the manuscript, and J. Erlich for help building the computing grid that all model fitting was conducted on.

## References

1. Machado A, Malheiro MT, Erlhagen W. Learning to Time: a perspective. J Exp Anal Behav. 2009;92: 423–458.

2. Mauk MD, Buonomano DV. The neural basis of temporal processing. Annu Rev Neurosci. 2004;27: 307–340.

3. Buhusi CV, Meck WH. What makes us tick? Functional and neural mechanisms of interval timing. Nat Rev Neurosci. 2005;6: 755–765.

4. Muller T, Nobre AC. Perceiving the passage of time: neural possibilities. Ann N Y Acad Sci. 2014;1326: 60–71.

5. Emmons EB, De Corte BJ, Kim Y, Parker KL, Matell MS, Narayanan NS. Rodent Medial Frontal Control of Temporal Processing in the Dorsomedial Striatum. J Neurosci. 2017;37: 8718–8733.

6. Merchant H, Harrington DL, Meck WH. Neural basis of the perception and estimation of time. Annu Rev Neurosci. 2013;36: 313–336.

7. Allman MJ, Teki S, Griffiths TD, Meck WH. Properties of the internal clock: first- and second-order principles of subjective time. Annu Rev Psychol. 2014;65: 743–771.

8. Gibbon J, Church RM. Time left: linear versus logarithmic subjective time. J Exp Psychol Anim Behav Process. 1981;7: 87–107.

9. Jozefowiez J, Machado A. On the content of learning in interval timing: representations or associations? Behav Processes. 2013;95: 8–17.

10. Yi L. Do rats represent time logarithmically or linearly? Behav Processes. 2009;81: 274–279.

11. Lake JI, LaBar KS, Meck WH. Emotional modulation of interval timing and time perception. Neurosci Biobehav Rev. 2016;64: 403–420.

12. Droit-Volet S. Time perception, emotions and mood disorders. J Physiol Paris. 2013;107: 255–264.

13. Wittmann M. The inner experience of time. Philos Trans R Soc Lond B Biol Sci. 2009;364: 1955–1967.

14. Wearden JH. Human performance on an analogue of an interval bisection task. Q J Exp Psychol B. 1991;43: 59–81.

15. Jozefowiez J, Staddon JER, Cerutti DT. The behavioral economics of choice and interval timing. Psychol Rev. 2009;116: 519–539.

16. Kopec CD, Brody CD. Human performance on the temporal bisection task. Brain Cogn. 2010;74: 262–272.

17. Ogden RS, Wearden JH, Jones LA. The remembrance of times past: interference in temporal reference memory. J Exp Psychol Hum Percept Perform. 2008;34: 1524–1544.

18. Spetch ML, Wilkie DM. A systematic bias in pigeons’ memory for food and light durations. Behaviour Analysis Letters. Elsevier Science Publishers, B.V.; 1982; Available: http://psycnet.apa.org/psycinfo/1983-12001-001

19. Eagleman DM. Human time perception and its illusions. Curr Opin Neurobiol. 2008;18: 131–136.

20. Buhusi CV, Meck WH. Relative time sharing: new findings and an extension of the resource allocation model of temporal processing. Philos Trans R Soc Lond B Biol Sci. 2009;364: 1875–1885.

21. Church RM, Deluty MZ. Bisection of temporal intervals. J Exp Psychol Anim Behav Process. 1977;3: 216–228.

22. Gibbon J. On the form and location of the Psychometric Bisection Function for time. J Math Psychol. 1981;24: 58–87.

23. de Carvalho MP, Machado A, Vasconcelos M. Animal timing: a synthetic approach. Anim Cogn. 2016;19: 707–732.

24. Allan LG, Gibbon J. Human bisection at the geometric mean. Learn Motiv. 1991;22: 39–58.

25. Wearden JH, Ferrara A. Stimulus range effects in temporal bisection by humans. Q J Exp Psychol B. 1996;49: 24–44.

26. Allan LG. The location and interpretation of the bisection point. Q J Exp Psychol B. 2002;55: 43–60.

27. Droit-Volet S. Alerting attention and time perception in children. J Exp Child Psychol. 2003;85: 372–384.

28. Droit-Volet S, Rattat A-C. A further analysis of time bisection behavior in children with and without reference memory: the similarity and the partition task. Acta Psychol. 2007;125: 240–256.

29. Stubbs DA. Scaling of stimulus duration by pigeons. J Exp Anal Behav. 1976;26: 15–25.

30. Platt JR, Davis ER. Bisection of temporal intervals by pigeons. J Exp Psychol Anim Behav Process. 1983;9: 160–170.

31. Meck WH. Selective adjustment of the speed of internal clock and memory processes. J Exp Psychol Anim Behav Process. 1983;9: 171–201.

32. Siegel SF. A test of the similarity rule model of temporal bisection. Learn Motiv. 1986;17: 59–75.

33. Killeen PR, Fetterman JG. A behavioral theory of timing. Psychol Rev. 1988;95: 274–295.

34. Penney TB, Gibbon J, Meck WH. Categorical scaling of duration bisection in pigeons (Columba livia), mice (Mus musculus), and humans (Homo sapiens). Psychol Sci. 2008;19: 1103–1109.

35. Vieira de Castro AC, Machado A, Tomanari GY. The context effect as interaction of temporal generalization gradients: testing the fundamental assumptions of the Learning-to-Time model. Behav Processes. 2013;95: 18–30.

36. Gibbon J. Scalar expectancy theory and Weber’s law in animal timing. Psychol Rev. 1977;84: 279–325.

37. Gibbon J, Fairhurst S. Ratio versus difference comparators in choice. J Exp Anal Behav. 1994;62: 409–434.

38. Buhusi CV, Aziz D, Winslow D, Carter RE, Swearingen JE, Buhusi MC. Interval timing accuracy and scalar timing in C57BL/6 mice. Behav Neurosci. 2009;123: 1102–1113.

39. Church RM, Meck WH, Gibbon J. Application of scalar timing theory to individual trials. J Exp Psychol Anim Behav Process. 1994;20: 135–155.

40. Fetterman JG, Killeen PR. Time discrimination in Columba livia and Homo sapiens. J Exp Psychol Anim Behav Process. 1992;18: 80–94.

41. Gibbon J, Church RM, Meck WH. Scalar timing in memory. Ann N Y Acad Sci. 1984;423: 52–77.

42. Lejeune H, Wearden JH. Scalar properties in animal timing: conformity and violations. Q J Exp Psychol. 2006;59: 1875–1908.

43. Rakitin BC, Gibbon J, Penney TB, Malapani C, Hinton SC, Meck WH. Scalar expectancy theory and peak-interval timing in humans. J Exp Psychol Anim Behav Process. 1998;24: 15–33.

44. Ainslie G, Herrnstein RJ. Preference reversal and delayed reinforcement. Anim Learn Behav. 1981;9: 476–482.

45. Green L, Myerson J, McFadden E. Rate of temporal discounting decreases with amount of reward. Mem Cognit. 1997;25: 715–723.

46. Killeen PR. An additive-utility model of delay discounting. Psychol Rev. 2009;116: 602–619.

47. Lane SD, Cherek DR, Pietras CJ, Tcheremissine OV. Measurement of delay discounting using trial-by-trial consequences. Behav Processes. 2003;64: 287–303.

48. Mazur JE. Tradeoffs among delay, rate, and amount of reinforcement. Behav Processes. 2000;49: 1–10.

49. Myerson J, Green L. Discounting of delayed rewards: Models of individual choice. J Exp Anal Behav. 1995;64: 263–276.

50. Peters J, Büchel C. Episodic future thinking reduces reward delay discounting through an enhancement of prefrontal-mediotemporal interactions. Neuron. 2010;66: 138–148.

51. Pine A, Shiner T, Seymour B, Dolan RJ. Dopamine, Time, and Impulsivity in Humans. Journal of Neuroscience. 2010;30: 8888–8896.

52. Rachlin H, Raineri A, Cross D. Subjective probability and delay. J Exp Anal Behav. 1991;55: 233–244.

53. Richards JB, Mitchell SH, de Wit H, Seiden LS. Determination of discount functions in rats with an adjusting-amount procedure. J Exp Anal Behav. 1997;67: 353–366.

54. Rodriguez ML, Logue AW. Adjusting delay to reinforcement: comparing choice in pigeons and humans. J Exp Psychol Anim Behav Process. 1988;14: 105–117.

55. Ratcliff R. A theory of memory retrieval. Psychol Rev. 1978;85: 59–108.

56. Tajima S, Drugowitsch J, Pouget A. Optimal policy for value-based decision-making. Nat Commun. 2016;7: 12400.

57. Krajbich I, Armel C, Rangel A. Visual fixations and the computation and comparison of value in simple choice. Nat Neurosci. 2010;13: 1292–1298.

58. Einstein A. Über die von der molekularkinetischen Theorie der Wärme geforderte Bewegung von in ruhenden Flüssigkeiten suspendierten Teilchen. Ann Phys. 1905;322: 549–560.

59. Wald A. On Cumulative Sums of Random Variables. Ann Math Stat. 1944;15: 283–296.

60. Schrödinger E. Zur theorie der fall-und steigversuche an teilchen mit brownscher bewegung. Phys Z. 1915;16: 289–295.

61. Green L, Myerson J, Holt DD, Slevin JR, Estle SJ. Discounting of delayed food rewards in pigeons and rats: is there a magnitude effect? J Exp Anal Behav. 2004;81: 39–50.

62. Roberts S. Isolation of an internal clock. J Exp Psychol Anim Behav Process. 1981;7: 242–268.

63. Church RM, Gibbon J. Temporal generalization. J Exp Psychol Anim Behav Process. 1982;8: 165–186.

64. Oliveira L, Machado A. Context effect in a temporal bisection task with the choice keys available during the sample. Behav Processes. 2009;81: 286–292.

65. Machado A. Learning the temporal dynamics of behavior. Psychol Rev. 1997;104: 241–265.

66. Stubbs A. The discrimination of stimulus duration by pigeons. J Exp Anal Behav. 1968;11: 223–238.

67. Pavlov IP, Anrep GV. Conditioned Reflexes. An Investigation of the Physiological Activity of the Cerebral Cortex… Translated and Edited by GV Anrep. London; 1927.

68. Machado A, Guilhardi P. Shifts in the psychometric function and their implications for models of timing. J Exp Anal Behav. 2000;74: 25–54.

69. Skinner BF. The Extinction of Chained Reflexes. Proc Natl Acad Sci USA. 1934;20: 234–237.

70. Guilhardi P, Macinnis MLM, Church RM, Machado A. Shifts in the psychophysical function in rats. Behav Processes. 2007;75: 167–175.

71. Gouvêa TS, Monteiro T, Soares S, Atallah BV, Paton JJ. Ongoing behavior predicts perceptual report of interval duration. Front Neurorobot. 2014;8: 10.

72. Matell MS, Meck WH. Cortico-striatal circuits and interval timing: coincidence detection of oscillatory processes. Brain Res Cogn Brain Res. 2004;21: 139–170.

73. Bakhurin KI, Goudar V, Shobe JL, Claar LD, Buonomano DV, Masmanidis SC. Differential Encoding of Time by Prefrontal and Striatal Network Dynamics. J Neurosci. 2017;37: 854–870.

74. Soares S, Atallah BV, Paton JJ. Midbrain dopamine neurons control judgment of time. Science. 2016;354: 1273–1277.

75. Finnerty GT, Shadlen MN, Jazayeri M, Nobre AC, Buonomano DV. Time in Cortical Circuits. J Neurosci. 2015;35: 13912–13916.

76. Cui X. Hyperbolic discounting emerges from the scalar property of interval timing. Front Integr Neurosci. 2011;5: 24.

77. Takahashi T. Time-estimation error following Weber-Fechner law may explain subadditive time-discounting. Med Hypotheses. 2006;67: 1372–1374.

78. Mazur JE, Coe D. Tests of transitivity in choices between fixed and variable reinforcer delays. J Exp Anal Behav. 1987;47: 287–297.

79. Roberts S. Cross-modal use of an internal clock. J Exp Psychol Anim Behav Process. 1982;8: 2–22.

80. Church RM, Broadbent HA. Alternative representations of time, number, and rate. Cognition. 1990;37: 55–81.

81. Lejeune H, Ferrara A, Soffíe M, Bronchart M, Wearden JH. Peak procedure performance in young adult and aged rats: acquisition and adaptation to a changing temporal criterion. Q J Exp Psychol B. 1998;51: 193–217.

82. Gallistel CR, King A, McDonald R. Sources of variability and systematic error in mouse timing behavior. J Exp Psychol Anim Behav Process. 2004;30: 3–16.

